# Mass photometric detection and quantification of nanoscale α-synuclein phase separation

**DOI:** 10.1101/2022.05.03.490467

**Authors:** Soumik Ray, Thomas O. Mason, Lars Boyens-Thiele, Nadin Jahnke, Alexander K. Buell

**Author notes:** Correspondence: Prof. Alexander K. Buell, Department of Biotechnology and Biomedicine, Technical University of Denmark, Søltofts Plads, Building 227, 2800, Kgs. Lyngby, Denmark.

## Abstract

α-Synuclein (α-Syn) liquid-liquid phase separation (LLPS) leads to irreversible amyloid fibril formation associated with Parkinson’s disease pathogenesis. Critical concentrations of α-Syn LLPS are relatively high under physiological solution conditions. Moreover, α-Syn exhibits delayed LLPS kinetics under certain conditions which deviates from the behaviour predicted by classical homogeneous nucleation theory. In the current body of work, using interferometric light scattering (iSCAT), also known as mass photometry, we experimentally probe that α-Syn can form nanoscale phase separated assemblies/clusters, containing tens to hundreds of molecules— both above and below the critical LLPS concentration down to physiologically relevant scales. The formation of these clusters is instantaneous, even under conditions where the formation of microscopically visible droplets takes several days. However, they account for a very small volume fraction below saturation concentration. The slow growth of the nanoclusters can be attributed to a kinetic barrier which can be overcome by increasing the solution temperature to just below the droplet melting point. We provide reasons for caution in quantifying dilute phase concentrations for α-Syn LLPS samples containing nanoscale droplets—which can only be separated using ultracentrifugation. In addition, we also delineate that the presence of certain surfaces facilitates α-Syn droplet nucleation under conditions of delayed kinetics but is not a mandatory prerequisite for nanocluster formation. Taken together, our findings reveal that phase separation of α-Syn occurs at a wider range of solution conditions than predicted so far and provides an important step towards understanding α-Syn LLPS within physiological scales.

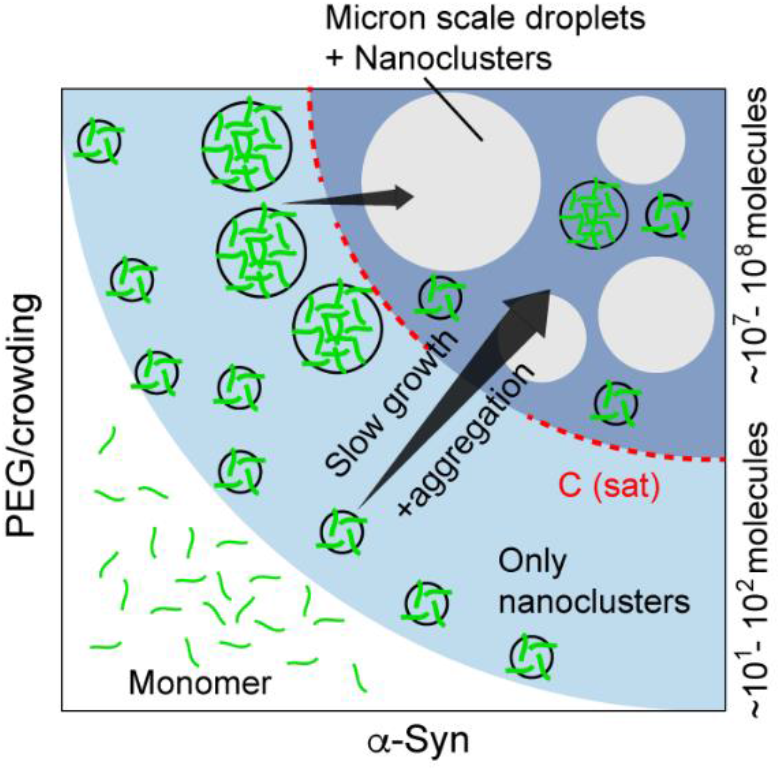

## Introduction

Biomolecular condensates have received increasing attention in recent years due to their widespread implications, in the form of membraneless organelles (MLOs), in health and disease^1–27^. MLOs are thought to form via a process called liquid-liquid phase separation (LLPS)^28–31^, in which homogeneous mixtures of molecules in an aqueous solution, under certain conditions, spontaneously demix into two (or more) phases: a dense phase (comprising of spherical droplets) and a dilute phase in the simplest case. Just after their formation, phase separated droplets show liquid-like properties such as fusion upon contact, surface wetting and dripping^1,29^. With time, the droplets grow via fusion and/or Ostwald ripening^29,32,33^. At a given condition (e.g., ionic strength, pH, temperature, presence of crowding), protein molecules can spontaneously form liquid droplets above a certain bulk concentration, which is termed the critical or saturation concentration (C_sat_)^34^. Conditions that change the extent of intermolecular interaction (defined by the Flory parameter, χ^28,29,34–36^), also change the value of C_sat_—which ultimately defines the binodal phase boundaries of a given protein. χ is a measure for the relative strength of protein-protein, protein-solvent and solvent-solvent interactions and a negative value of χ signifies that protein-solvent interactions are favorable. If a protein solution is quenched into the binodal region of the phase diagram, the protein molecules are thought to undergo LLPS according to classical homogeneous nucleation theory^37–39^. In this theory, the nucleation and spontaneous growth of the initial droplets is governed by the free energy (ΔG) of a droplet which, for a given value of χ, depends on its size. The size-dependence of the free energy of droplet formation leads to a free energy barrier for droplet nucleation. This barrier is defined by the difference in the chemical potential between the dense and the dilute phase molecules (Δ*μ*), the interfacial tension of the droplets (γ), the size of the droplets (radius *r*), and can be described by the following equation:

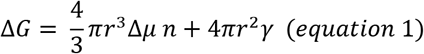

Where n is the number of molecules per unit volume. The two parts of the equation describe the volume energy and the surface energy, respectively. There is a competition between these two energies, which, depending on the radius (*r*) of the droplets, determines whether they will further grow or dissolve^34,37–39^. The surface energy always opposes phase separation. On the other hand, depending on χ and hence, Δ*μ*, the volume energy can be negative and can overcome the interfacial penalty above a certain number of molecules/droplet radius, leading to the growth of the new phase.

Protein LLPS is largely driven by transient, multivalent, intermolecular interactions which are defined by the interplay of amino acid sequence and solution conditions^40^. Intrinsically disordered regions (IDRs) and low complexity domains (LCDs) comprising of ‘sticker’ and ‘spacer’ motifs^41,42^ not only play a significant role in droplet formation but can also be used to tune the material properties of LLPS^43–50^. Intrinsically disordered proteins/peptides (IDPs) are also generally more susceptible towards LLPS compared to folded proteins. Interestingly, due to their tendency to misfold, there is an inherent risk associated with concentrating IDPs inside phase separated assemblies—aberrant, liquid-to-solid phase transition and toxic amyloid fibril formation^14,35,45,51–67^. The formation of amyloid fibrils does not necessarily require that the protein undergoes LLPS first, as there are often multiple pathways that lead from a homogeneous protein solution to amyloid fibrils. However, the highly concentrated environment inside dense condensate droplets strongly facilitates the formation of amyloid fibrils compared to the more dilute homogeneous solution scenario. In particular α-synuclein (α-Syn)^68^, which can misfold and aggregate into cytotoxic oligomers and amyloid fibrils associated with Parkinson’s disease (PD) and other synucleinopathies^69^ was until recently believed to form amyloid fibrils exclusively through heterogeneous nucleation on surfaces^70–73^. However, it is now becoming clear that α-Syn forms amyloid fibrils efficiently inside condensate droplets. α-Syn can undergo LLPS at physiological pH in the presence of molecular crowding induced by polyethylene glycol (PEG)^58^ or Ficoll^63^, however usually only at relatively high (>100 μM) concentrations. Moreover, under certain conditions, α-Syn LLPS can take hours to days to be visually confirmed by microscopy^58,60^. This phenomenon of delayed LLPS is most common for α-Syn, but not limited to it. The microtubule binding protein tau, associated with Alzheimer’s disease (AD) pathology, also has been reported to undergo delayed phase separation^53^.

These observations raise the question whether the bulk protein concentrations where delayed LLPS is observed is indeed inside the metastable region of the binodal (>C_sat_), as the low driving force for LLPS is indicative for a very small degree of supersaturation. It is also unclear whether the delay is due to a large nucleation barrier for the first stable droplets to form or, whether the growth of microscopically invisible protein assemblies into microscopically visible ones is the limiting factor. In this line, recent development in the field suggests that nanoscale protein clusters (e.g. of the fused in sarcoma, FUS protein) can form even in subsaturated concentrations (below C_sat_) where no macroscopic LLPS is observed^38^. Formation of such assemblies is hypothesized to help cross the nucleation barrier of LLPS above C_sat_. Nanoscale protein assembly persists even above C_sat_ (along with LLPS) for another well-studied phase separating protein, hnRNPA1^37^. Intriguingly, in the nanoscale regime, the growth behavior of these clusters deviates from that predicted by classical nucleation theory^37^.

In the present study, we experimentally demonstrate that α-Syn forms liquid-like, nanoscale clusters containing tens to hundreds of molecules at physiologically relevant concentrations (one order of magnitude below its typical saturation concentration). With increasing bulk protein concentration, the apparent mass and volume fraction of these nanoclusters increases predictably, and they persist even above C_sat_ along with larger droplets. Interestingly, factors that are known to modulate α-Syn LLPS also effects the nanoscale cluster formation in similar ways. We probe that the nucleation of α-Syn nanoclusters is spontaneous, however, their growth can be exceedingly slow under some conditions, which explains their delayed appearance as judged by state-of-the-art microscopic techniques^58^. We find that the slow growth is likely due to a high kinetic barrier that prevents the nanoclusters from growing into larger droplets. This barrier can be overcome by increasing the solution temperature to values just below the dissolution temperature of the droplets. Lastly, we delineate that although α-Syn droplets can nucleate on surfaces, nanocluster and subsequent amyloid fibril formation does not require surfaces. Taken together, our study provides experimental evidence of the two different size regimes (nanoscale and microscale) of α-Syn phase separated assemblies which has crucial physiological and pathological relevance. Our findings also give reason for caution in the use of centrifugation methods^74^ to elucidate protein phase diagrams and effects of surfaces in nucleating condensed states for systems that are able to undergo nanoscale phase separation^38,75,76^.

## Results

### Demixed α-Syn assemblies at different regions of its LLPS regime

To delineate the canonical LLPS regime of α-Syn under physiologically relevant solution conditions (20 mM PBS, pH 7.4, 25°C), we performed an orthogonal screen of α-Syn concentration and (%w/v) PEG-8000. Under these conditions, α-Syn spontaneously formed micron sized spherical phase separated assemblies above 100 μM in the presence of ≥ 15% (w/v) PEG-8000 (Figure 1a, left panel). The partitioning of 100 nM of Alexa-488 maleimide labeled 140C-α-Syn (Alexa488-140C-α-Syn) into the droplets confirmed that these assemblies indeed were protein-rich (Figure 1a, right panel). Just after their formation (day 0), α-Syn molecules showed high translational dynamics inside the droplets and recovered fully after photobleaching (FRAP) experiments (Figure 1b). The droplets also showed occasional fusion events suggestive of their liquid-like behavior (Figure 1b, inset). Interestingly, upon ageing, α-Syn liquid droplets underwent aberrant, liquid-to-solid phase transition. After 2 days of incubation at 25°C, the droplets became Thioflavin-T (ThT) positive—indicating the presence of amyloid fibrils (Figure 1c). Liquid-to-solid phase transition was further supported by the lack of fluorescence recovery in FRAP experiments at day 2 (Figure 1d), and bulk ThT -fluorescence based aggregation kinetics, where we observed that the ThT fluorescence increased steeply after 12-16h of incubation and reached a plateau after 1 day (Figure 1e). In order to check whether the increase in ThT fluorescence observed in this bulk experiment was associated with amyloid aggregation *inside* the droplets, we microscopically examined one α-Syn sample that had undergone LLPS (spiked with 50 μM ThT) for 48h with a cyan fluorescence channel (excitation 436 nm, emission 485 nm). We observed a progressive increase of ThT fluorescence inside individual droplets confirming that α-Syn LLPS indeed precedes its amyloid aggregation (Figure 1e, inset, Supplementary movie 1)^58^. Furthermore, in transmission electron microscopy (TEM) imaging, α-Syn droplets showed the presence of aggregate-like structures at day 4. The same sample showed the presence of amyloid fibrils in solution after 7-10 days of incubation (Figure 1f). All the experiments described in Figure 1b-f have been carried out using 200 μM α-Syn in presence of 20% (w/v) PEG-8000.

**Figure 1.**
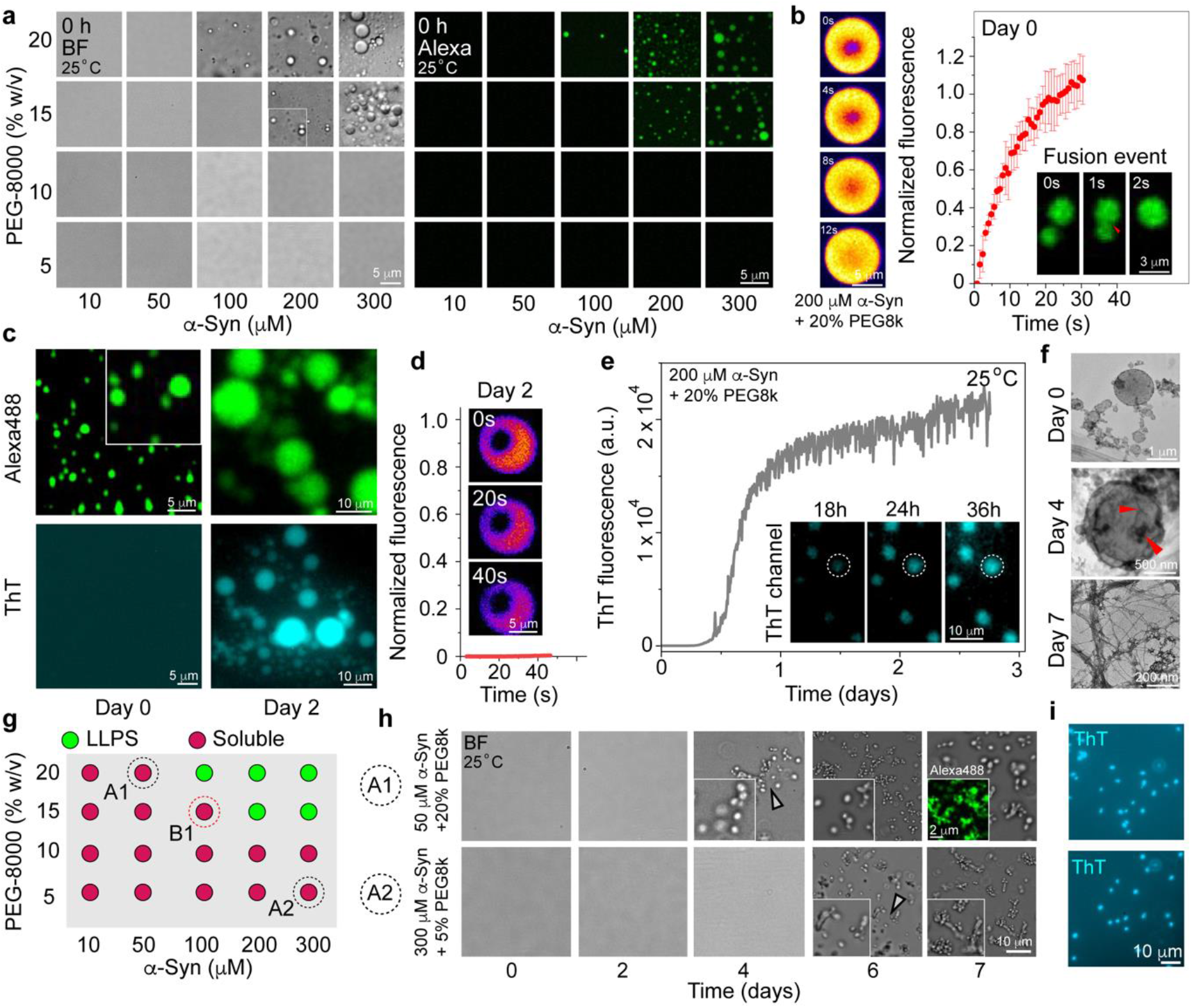
Characterization of α-Syn LLPS: **a**. (left panel) Representative bright field (BF) microscopy images of the LLPS regime of α-Syn (at 0h) in the presence of PEG-8000 at 25°C are shown. (Right panel) Representative fluorescence microscopy images of the LLPS regime of α-Syn (at 0h) spiked with 100 nM Alexa-488-140C-α-Syn at 25°C are shown. The experiments are performed 3 independent times with similar observation. **b**. (left panel) Representative snapshots of a α-Syn liquid droplet at different times post-bleaching. The images are shown in thermal lookup table (LUT) for better visualization. (Right panel) Normalized fluorescence recovery after photobleaching (FRAP) of α-Syn liquid droplets at 0h is shown. Values represent mean ± S.D for n=3 independent experiments. (Right panel, inset) Time-lapse fluorescence images showing 2 α-Syn liquid droplets fusing upon contact over the course of 2 s. **c**. Representative fluorescence images of α-Syn droplets (spiked with either Alexa488 or ThT) at day 0 and day 2 are shown under 2 fluorescence channels (Alexa488 (top) and CFP (bottom)). The experiment is performed 2 independent times with similar observations. **d**. Normalized fluorescence recovery after photobleaching (FRAP) of α-Syn droplets after 2 days is shown. (Inset) Representative snapshots of a α-Syn droplet at different times postbleaching. The images are shown in thermal LUT for better visualization. The experiment is performed 2 independent times with similar results. **e**. ThT kinetics of amyloid fibril formation is shown for a α-Syn LLPS system at 25°C. (Inset) Time-lapse images showing evolution of ThT fluorescence inside individual α-Syn droplets over the course of 36h. The experiment is performed 2 times with similar results. **f**. Representative TEM images of α-Syn LLPS solution at day 0, 4 and 7 are shown. Aggregated structures are denoted with red triangular pointers inside the day 4 droplets. The experiment is performed 2 times with similar observations. **g**. A schematic depicting the LLPS regime of α-Syn in the presence of PEG-8000. The green circles indicate LLPS (at 0h). The dotted circles (A1 and A2) represent conditions where clusters form after 4-6 days of incubation at 25°C. The dotted circle (B1) represents the condition where LLPS is observed after 1 day of incubation at 25°C (Supplementary Figure 1). **h**. Representative BF microscopy images of the conditions: A1 (50 μM α-Syn+20% (w/v) PEG-8000) and A2 (300 μM α-Syn+5% (w/v) PEG-8000), at different timepoints (day 0-7) are shown. The α-Syn clusters are denoted with white triangular pointers. The insets represent magnified images for better visualization. (Top right inset) Representative fluorescence microscopy image of the clusters under condition A1 at day 7, spiked with 100 nM of Alexa488-140C-α-Syn as label. **i**. Representative fluorescence images of α-Syn clusters spiked 50 μM ThT at day 7 are shown under CFP fluorescence channel. The experiments (**h-i**) are performed 3 independent times with similar observations. All the experiments described in Figure **1b-f** has been carried out using 200 μM α-Syn in presence of 20% (w/v) PEG-8000 in 20 mM PBS, pH 7.4.

Overall, our experiments provided a comprehensive characterization of α-Syn LLPS at physiological pH and at room temperature (25°C) (Figure 1a-g). Surprisingly, under certain conditions, the droplet state of α-Syn can take hours or even days to be visually confirmed by conventional optical microscopy techniques^58^. In order to check whether α-Syn can undergo delayed phase separation at conditions away from the canonical LLPS regime and below its apparent C_sat_ (termed as C_app_ in this manuscript), we incubated 50 μM and 300 μM α-Syn at 25°C in presence of 20% and 5% (w/v) PEG-8000, respectively (Figure 1g-h). Strikingly, we observed that for both conditions, distinct, protein-rich assemblies (several microns in size) started to appear after 4-6 days of incubation (Figure 1h, left panel, Supplementary movie 2). For now, we term these assemblies ‘clusters’, rather than droplets, because of their lack of liquid-like character and because we do not know whether they form through LLPS. Interestingly, the clusters were morphologically different from aggregates formed by concentrated monomeric α-Syn (700 μM, no PEG) in solution (Supplementary Figure 1, Supplementary movie 3) and showed solid-like behavior as soon as they were microscopically identified (day 4-6) (Supplementary Figure 1). They fluoresced strongly upon ThT addition (Figure 1i) and after 15 days of incubation formed higher-order structures containing amyloid fibrils as confirmed by TEM imaging (Supplementary Figure 1).

In another region of the phase diagram, as one approaches (but does not enter) the instantaneous LLPS regime (100 μM α-Syn + 15% (w/v) PEG-8000); α-Syn still showed spherical droplet formation, but with a delayed kinetics. Droplets appeared after 1 day of incubation at 25°C (Supplementary Figure 1) and the newly formed droplets showed ~80% recovery of fluorescence after photobleaching (Supplementary Figure 1), confirming their at least partially liquid-like character. Under these conditions, ThT fluorescence (amyloid aggregation) was only observed when the droplets were aged for at least 2 days (Supplementary Figure 1).

Our observations deviate from the commonly observed LLPS behavior where protein molecules spontaneously partition themselves (within seconds to minutes) into dense and dilute phases at concentrations above C_sat_. The question is whether the α-Syn clusters/droplets that form after hours/days are governed by the same types of interactions that lead to rapid LLPS under other conditions, and whether this delayed assembly is compatible with the prevailing view of protein phase separation. From a thermodynamic perspective, the delayed formation of α-Syn clusters/droplets away from its LLPS regime could be attributed to (***a***) A large nucleation energy barrier of assembly followed by rapid growth and maturation/phase transition (Supplementary Figure 1) and (***b***) A relatively smaller nucleation energy barrier followed by very slow growth by monomer incorporation and/or fusion (Supplementary Figure 1). In the second scenario (***b***), the first α-Syn assemblies could be both extremely small and sparse and the dilute phase monomers would likely dominate the signal in almost every in-solution spectroscopic technique (such as NMR), single molecule fluorescence measurements, and even traditional light scattering methods.

### Direct evidence of spontaneous α-Syn assembly below saturation concentration

The challenge was to experimentally probe whether microscopically invisible, higher order α-Syn assemblies can spontaneously form even under conditions where macroscopic LLPS is not observed. We hypothesized that interferometric light scattering (iSCAT), also known as mass photometry (MP) could be a way to probe the presence of α-Syn assemblies below C_app_^77–79^. This revolutionary new method allows accurate mass measurement of protein molecules and complexes in solution in the range of 40 kDa to 5 MDa—without the need of any labels (Figure 2a)^77^. The method is based on interferometric scattering to detect sub-wavelength particles with great precision and at single-molecule resolution. Moreover, MP is especially useful for α-Syn assemblies (with PEG-8000) because the detection limit of the instrument renders the α-Syn monomers (14 kDa) and PEG (8 kDa) molecules invisible. Additionally, in MP, the scattering intensity scales with the third power of the radius of a particle in solution^79^—making it resolve smaller protein assemblies more efficiently compared to other scattering-based methods. In MP measurements, the protein solution is drop-casted on a glass slide and a 525 nm laser illuminates the sample. When a protein molecule (or assembly of protein molecules) gets very close to the surface, the scattered light from the molecule(s) negatively interferes with the incident light reflected from the glass slide surface—creating a dark spot on a grey background (Figure 2a), which can be fitted with a Gaussian function. The peak height of the Gaussian fit (the most abundant contrast) linearly correlates with the average mass of the molecule(s) from a standard curve.

**Figure 2.**
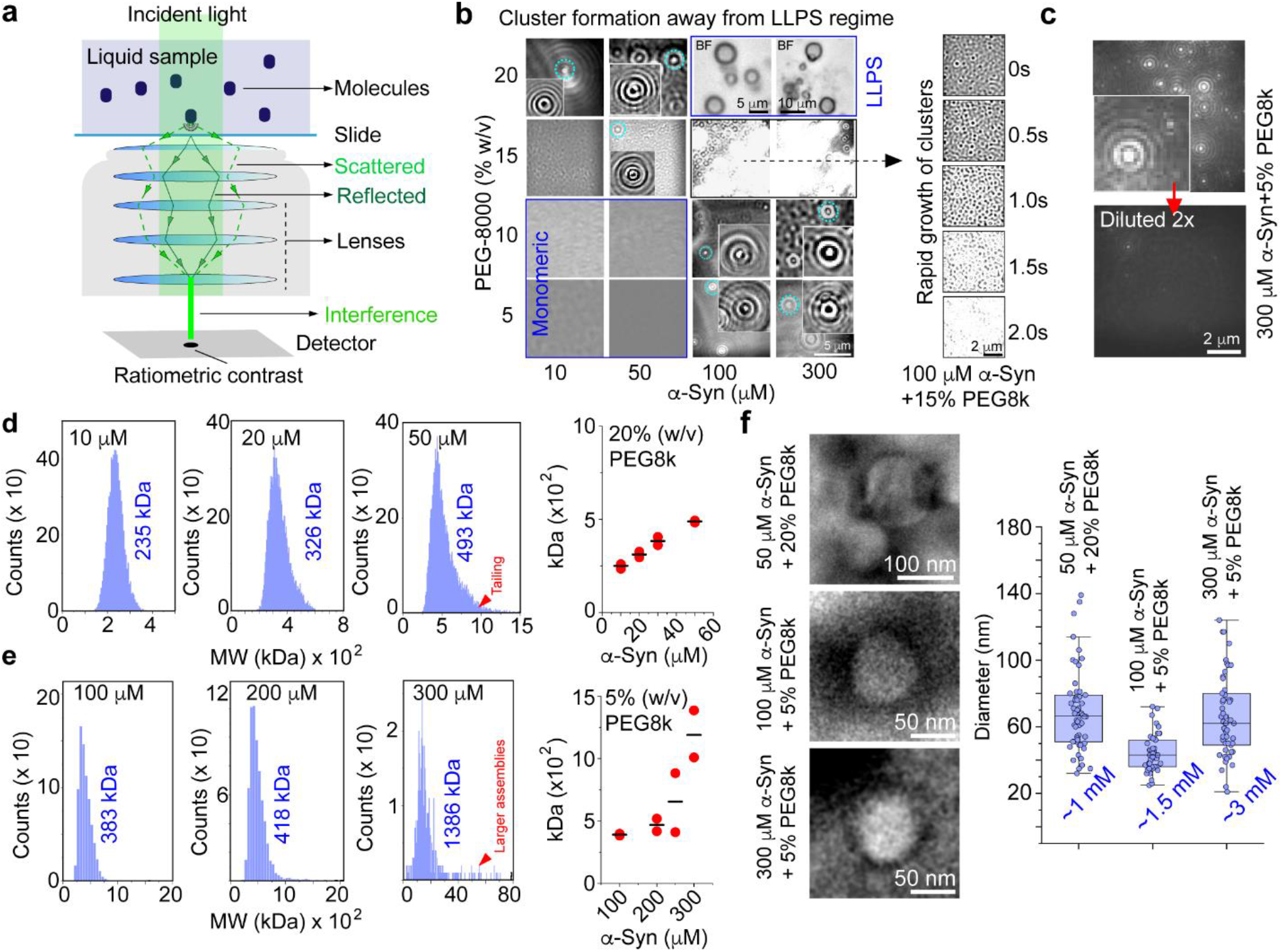
Characterization of α-Syn nanoclusters below Capp: **a**. Schematic describing the working principle behind the mass photometry (MP) instrument. An incident laser (525 nm) is used to illuminate the protein sample on a glass slide. Very close to the surface of the slide, the scattered light from the molecules/assemblies and the reflected light can interfere destructively, creating a dark spot on a grey background. The ratiometric contrast of the dark spot directly correlates with the mass/size of the molecules/assemblies. **b**. (left panel) The LLPS regime of α-Syn (0h) as seen in MP. The α-Syn clusters below C_app_ are marked with dashed circles (cyan) and shown as insets (ratiometric channel) for better visualization. The monomeric and LLPS regime is marked with blue rectangles. (Right panel) MP images (native channel) of a sample very close to the LLPS regime (100 μM α-Syn+15% (w/v) PEG-8000) is shown at different timepoints from 0-2 s. The experiment is performed 2 independent times with similar observations. **c**. MP images (native channel) of 300 μM α-Syn+5% (w/v) PEG-8000 before (top) and after (bottom) 2X dilution with buffer are shown. The experiment is performed 2 independent times with similar observations. **d-e**. (left panels) Molecular weight (MW) histograms obtained from MP measurements of 10, 20 and 50 μM α-Syn in presence of 20% (w/v) PEG-8000 (**d**) and 100, 200 and 300 μM α-Syn in presence of 5% (w/v) PEG-8000 (**e**) are shown. The average MW is noted in blue. The red triangular pointers (50 and 300 μM) denote tailing of the histogram due to presence of larger assemblies and/or surface wetting. (Right panels) The MW of the clusters are plotted against the total protein concentrations. Values are reported for n=2 independent experiments. The mean is denoted as a black dash. **f**. Representative TEM images of 50 μM α-Syn + 20% (w/v) PEG-8000, 100 μM α-Syn + 5% (w/v) PEG-8000 and 300 μM α-Syn + 5% (w/v) PEG-8000 showing presence of nanoscale spherical assemblies. **g**. The size distribution (diameter in nm) of α-Syn nanoclusters under different conditions is reported as box-whisker plots for N=50 clusters for each sample. Estimates of the maximum apparent concentration of α-Syn inside the nanoclusters are reported in blue.

We characterized various α-Syn+PEG-8000 concentration pairs by MP (Figure 2b, left panel, Supplementary Figure 2). To our surprise, our data showed that immediately after addition of PEG, α-Syn spontaneously assembled into higher order assemblies (clusters) even at physiologically relevant, low μM concentrations (one order of magnitude lower concentrations than those at which we observe droplets by conventional microscopy) (Figure 2b, left panel, Supplementary Figure 2, Supplementary movie 4). At very low protein and PEG concentrations, we did not detect any supramolecular assemblies—indicating that similar to visible droplet formation, these clusters also had a defined existence regime (Figure 2b, left panel, Supplementary Figure 2). Interestingly, very close to the LLPS regime (100 μM α-Syn+15% (w/v) PEG-8000), MP showed a rapid growth of the clusters—eventually saturating the detector within a few seconds (Figure 2b, right panel, Supplementary movie 5). We also observed that the clusters could be readily dissolved upon dilution (Figure 2c) and showed surface wetting (Supplementary movie 6)—both suggesting liquid-like properties similar to those of droplets formed by LLPS. Next, we used a set of standard protein samples of known molecular weights (BSA (66 kDa), urease (91 kDa, monomer) and thyroglobulin (669 kDa)) to generate a standard curve from their ratiometric contrasts (Supplementary Figure 2). From the standard curve calibration, we found that the apparent mass of the α-Syn clusters ranged from as low as 235 kDa (~16 α-Syn molecules) to 1386 kDa (~100 α-Syn molecules) depending on the protein and PEG concentration (Figure 2e, left panels). Intriguingly, the apparent mass of the clusters increased with increasing protein concentrations—reflecting more and more individual binding events below Capp before microscopically visible phase separation occurred (Figure 2d-e, right panels). However, closer to Capp, we often observed a tailing in the histogram and appearance of many larger assemblies outside the main Gaussian peak (Figure 2d-e, left panels, Supplementary Figure 2). For high α-Syn concentrations, the tailing effect was in the range of ~4000-12000 kDa (~300-900 α-Syn molecules)—which is beyond the size analysis limit of MP, and therefore these mass values are likely to be only very approximate. This could be due to presence of very large assemblies, but also could be a result of extensive surface wetting (Figure 2d-e, left panels, Supplementary Figure 2). To note, these clusters could also form in the presence of PEG-3000 and Ficoll-400 (two other crowding agents)—indicating that generic molecular crowding was sufficient to induce their formation (Supplementary Figure 2). It is important to note that at high protein concentrations, there could be a ‘low mass noise’ due to fluctuations of protein densities on the surface. Therefore, the MW values of the α-Syn clusters are possibly not absolute, rather a close approximate.

High NaCl concentrations and low pH promote α-Syn LLPS^58,59^. We wanted to probe whether this dependence was also observed for cluster formation. Our data showed that in the presence of 250 mM NaCl (20 mM PBS buffer, pH 7.4), α-Syn readily formed clusters with sizes >1000 kDa (>70 α-Syn molecules)—which was substantially compromised in an identical sample which was prepared without NaCl (20 mM PB buffer, pH 7.4) (Supplementary Figure 2). Furthermore, titrating small volumes of concentrated HCl (50 mM) into 20 μM α-Syn+10% (w/v) PEG-8000 (pH 7.4) led to formation of α-Syn clusters containing ~50 α-Syn molecules (712 kDa) when the solution pH was lowered to ~6.5 (Supplementary Figure 2).

Subsequently, we employed TEM imaging to gather visual evidence of α-Syn clusters to quantify their approximate dimensions and hence enable an estimate of the protein concentration inside the clusters. We chose 50 μM α-Syn + 20% (w/v) PEG-8000; 100 and 300 μM α-Syn + 5% (w/v) PEG-8000 for our TEM studies. Strikingly, all three samples showed the presence of nano-scale spherical assemblies (Figure 2f, left panel) with an average diameter of ~40-60 nm (Figure 2f, right panel). TEM imaging combined with MP measurements provided us with very useful information: (***1***) the approximate volume of the spherical nanoscale-clusters (nanoclusters) and (***2***) the corresponding apparent mass (molecule number) of the clusters. We considered the minimum volume of the clusters from the size distribution data (Figure 2f, right panel) and calculated the maximum apparent concentration of α-Syn to be ~1.0-3.0 mM inside these nanoclusters (Figure 2f, right panel).

To further strengthen our observations, we took different concentrations of α-Syn and PEG-8000 and subjected them to dynamic light scattering (DLS) experiments. In DLS of sub-wavelength particles, the scattering intensity scales with the sixth power of the radius of a particle in a solution (Rayleigh scattering)—and therefore DLS is a useful tool for identifying very low concentrations of higher order protein assemblies even with an excess of background monomer. Monomeric α-Syn (100 μM) showed a single DLS peak at ~3.2 nm, which agrees well with the known hydrodynamic radius of full-length α-Syn ^80,81^(Supplementary Figure 3). The amplitude and shape of the autocorrelation function suggested a homogeneous distribution of sizes in the monomeric α-Syn solution (Supplementary Figure 3). Next, we analyzed the entire α-Syn nanocluster and LLPS regime (at pH 7.4) using DLS. Intriguingly, we observed that the relative monomer peak intensity progressively decreased with increasing protein concentrations (at constant PEG-8000 (5-20% (w/v)), along with the presence of multiple, low intensity peaks larger than 3.4 nm (Supplementary Figure 3). In addition, the amplitude of the autocorrelation function was substantially increased for Tau>1E-05 s, indicating formation of higher order assemblies (Supplementary Figure 3) at high protein and/or PEG concentrations (Supplementary Figure 3). These effects were most prominent above C_app_ (LLPS)— however, some indication of the presence of species other than monomeric α-Syn was also observed under conditions where we observed nanoclusters in MP. To note, PEG-8000 alone did not show any DLS signal at 20% (w/v) concentration (maximum concentration used in our experiments) (Supplementary Figure 3). Importantly, as one increases the PEG-8000 concentration (and therefore, viscosity), the monomer peak is expected be shifted towards a higher apparent size unless a viscosity correction is applied. On the contrary, we often observed an overall shift to a lower apparent size of the monomeric peak in the samples under conditions where larger assemblies existed (in presence of PEG)—possibly due to limitations in the fitting of DLS signals stemming from very heterogeneous solutions.

### Temporal evolution of α-Syn nanoclusters

From our previous experiments, we know that micron-sized assemblies below Capp can be visually identified using conventional microscopy when the solutions are aged for 4-6 days (Figure 1h) at 25°C. We have also established that the formation of the nanoclusters is rapid, i.e., they form within the time scale (seconds) of our MP experiment (Figure 2, Supplementary Figure 2, Supplementary Figure 1-scenario ***b***). If nanoclusters arise from LLPS, their temporal evolution may be explained by two mechanisms: (***I***) under conducive solution conditions (inside the binodal), all nanoclusters spontaneously form immediately, and the volume fractions remain the same even after 4-6 days. This means that the cluster growth by fusion/ripening is very slow (Figure 3a). This hypothesis also demands that the nanoclusters remain liquid-like for a considerable amount of time to allow efficient growth. Or (***II***) very few nanoclusters form at the beginning and they gradually incorporate monomer from the solution, changing the dilute and dense phase volume fractions progressively (Figure 3a). In the second scenario, it is also possible that new nucleation events occur throughout the incubation period contributing to a continuous change in volume fraction.

**Figure 3.**
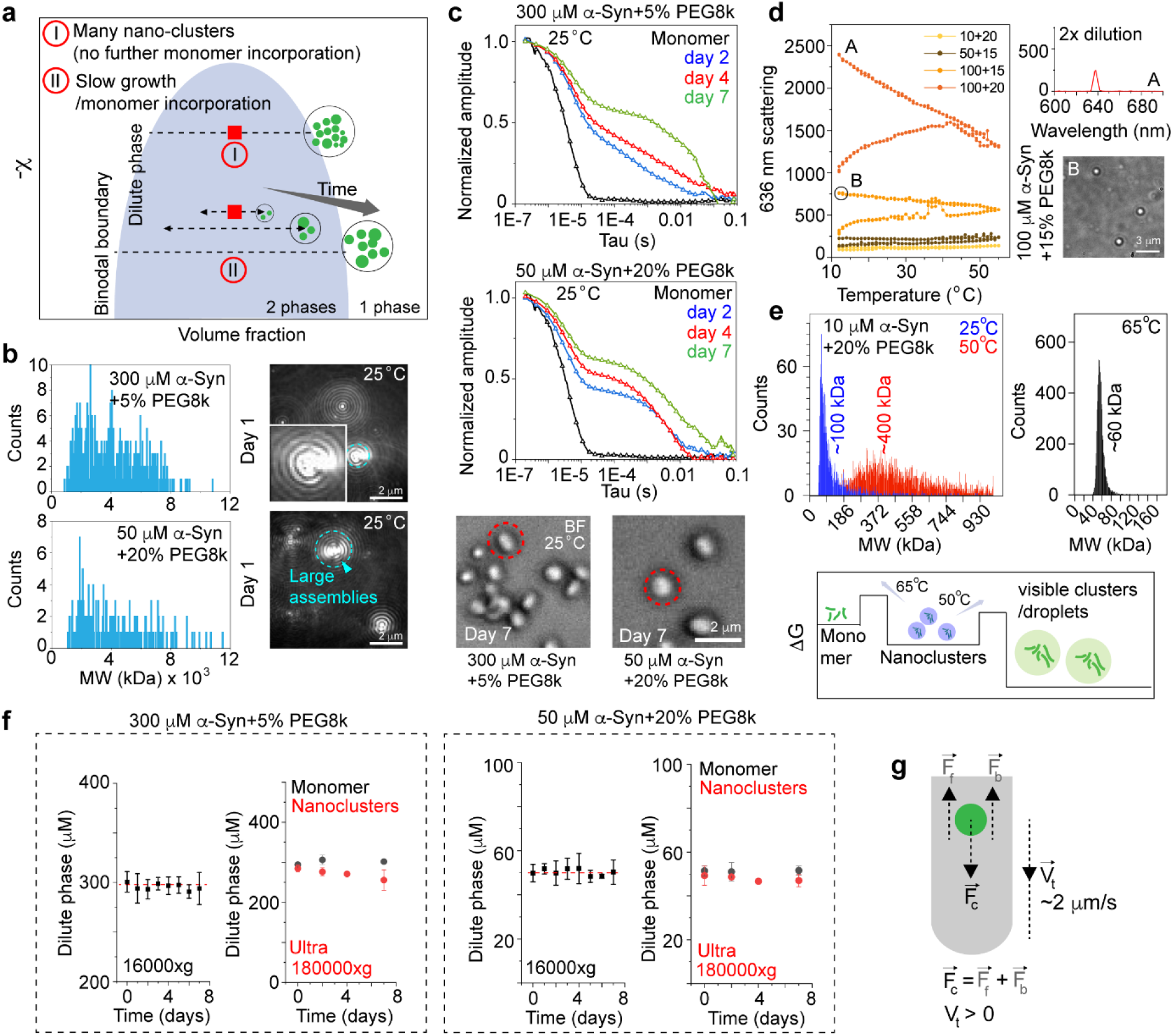
Mechanism of growth for α-Syn nanoclusters into micron-scale assemblies: **a**. Schematic explaining two possible scenarios which could explain the delayed growth of α-Syn nanoclusters assuming that they are, indeed, phase separated assemblies. In scenario (I), all the nanoclusters form spontaneously upon addition of PEG. However, their growth via fusion/ripening is very slow. In scenario (II), new nanoclusters form throughout the incubation period. However, they slowly incorporate more and more monomer from the solution as they grow—changing the volume fractions constantly. The X axis of the schematic represents the volume fraction, and the Y axis represents the χ parameter of LLPS^36^. The blue area represents the binodal and the red squares represents the nanocluster forming solution conditions. **b**. (left panels) MP molecular weight (MW) analysis of 300 μM α-Syn+5% (w/v) PEG-8000 (top) and 50 μM α-Syn+20% (w/v) PEG-8000 (bottom) after 1 day of incubation at 25°C. (Right panels) Corresponding MP images (native channel) showing the existence of large assemblies in the solutions (marked with cyan dashed circles and provided as an inset for better visualization). The experiment is performed twice with similar results. **c**. (upper panel) The normalized amplitude of the autocorrelation function (obtained in DLS) for 300 μM α-Syn+5% (w/v) PEG-8000 (top) and 50 μM α-Syn+20% (w/v) PEG-8000 (bottom) at day 2, 4 and 7 are shown. The autocorrelation function obtained for monomeric 100 μM α-Syn is also plotted as a reference in the same graphs (in black) for comparison. The X-axis (Tau) is reported in log scale. The experiment is performed 3 times with similar results. (Lower panel) Representative BF microscopy images showing the presence of α-Syn clusters in the same samples after 7 days of incubation at 25°C. The red dashed circles denote individual α-Syn clusters. **d**. (left panel) Static light scattering (SLS) intensity (636 nm) of 100 μM α-Syn+20% (w/v) PEG-8000, 100 μM α-Syn+15% (w/v) PEG-8000, 50 μM α-Syn+15% (w/v) PEG-8000 and 10 μM α-Syn+20% (w/v) PEG-8000 (prepared in 20 mM PBS, pH 7.4) is shown while the sample is heated from 12°C to 55°C and back to 12°C. (Right panel) After completion of the scattering experiment (at point A), the sample is diluted 2 times with buffer and the scattering intensity (636 nm) is plotted at 12°C. The bottom panel shows a representative BF microscopy image of the droplets after completion of the SLS experiment (at point B). The experiments are performed 2 times with similar results. **e**. MW histograms obtained from MP measurements of 10 μM α-Syn+20% (w/v) PEG-8000 at different temperatures (25°C, 50°C and 65°C) are shown. (Lower panel) Schematic describing a possible energy landscape of α-Syn nanoclusters with exceedingly slow growth rates. **f**. Concentration measurements of the dilute phase at different time-points (day 0-7) after centrifuging the nanocluster forming samples at 16000Xg (normal centrifugation) (left) and at 180000Xg (ultracentrifugation) (right). The ultracentrifugation data also show dilute phase concentrations of the monomer controls (in black) where no nanoclusters are present. Values represent mean ±S.D. for n=3 independent experiments. **g**. Schematic representation of the mathematical interpretation used in our study to check whether the nanoclusters can be ultracentrifuged at 180000Xg.

To probe the growth of α-Syn nanoclusters, we employed MP and DLS studies with 50 and 300 μM α-Syn in the presence of 20 and 5% (w/v) PEG-8000, respectively in a time dependent manner. In MP, we observed that the apparent mass of the nanoclusters increased significantly after 1 day of incubation at 25°C (1000-4000 kDa). Due to the presence of many large assemblies, the detector was saturated, and the instrument could no longer quantify the average mass (Figure 3b). This phenomenon was also evident in DLS, where we observed a progressive increase in the amplitude of the autocorrelation function with time (Figure 3c), along with appearance of new signals at higher hydrodynamic radius (Supplementary Figure 4). Monomeric α-Syn (100 μM) and 50 μM α-Syn+5% (w/v) PEG-8000 where no nanoclusters formed, showed minimal change in DLS signal and their respective autocorrelation functions even after 7 days of incubation at 25°C (Supplementary Figure 4). We also noted a systematic decrease in the relative area of the monomer peak (1-10 nm) with time for 300 μM α-Syn+5% (w/v) PEG-8000. Although a decrease in the peak area could not be used directly to quantify the concentration/amount of the monomer, it reliably indicates the formation of larger assemblies over time (Supplementary Figure 4). The monomer peak area remained constant under conditions where no clusters formed (Supplementary Figure 4). α-Syn nanoclusters induced by PEG-3000 were also characterized by DLS during their growth. Our data showed that along with a decreased peak intensity, the monomer peak of α-Syn was substantially widened with an increase in the amplitude of the autocorrelation function after 2 days of incubation at 25°C (Supplementary Figure 5). To note, monomer peak widening was more prominently observed for PEG-3000 compared to PEG-8000, possibly because PEG-3000 induced assemblies were very close to the size of α-Syn monomers.

Unlike most protein LLPS systems, α-Syn droplets persist upon heating in the temperature range of 12-55°C ^60^. However, we found that upon prolonged incubation at 65°C, droplets formed by 100 μM α-Syn+20% (w/v) PEG-8000 could indeed be dissolved^58^ as evident from the decrease in the cumulant hydrodynamic radius in DLS measurements to ~3 nm (monomeric) (Supplementary Figure 6). As liquid-to-solid transition progressed (0h-48h), droplets became more and more resistant towards heating (Supplementary Figure 6). To check whether α-Syn droplets under milder LLPS conditions (200 μM α-Syn+20% (w/v) PEG-8000 prepared in 10 mM PB, 40 mM NaCl, pH 7.4) were more susceptible towards heat dissolution, we employed static light scattering (SLS) (636 nm) measurements in the temperature range of 12-55°C. In this experiment, we first increased the solution temperature from 12°C to 55°C and reverted to 12°C subsequently. To our surprise, we observed that after the first heating step (12→55°C), the SLS intensity was substantially increased (with appearance of many new droplets) when the solution was subsequently cooled down to 12°C (Supplementary Figure 6). This phenomenon was also observed for another LLPS condition (100 μM α-Syn+20% (w/v) PEG-8000 in 20 mM PBS, pH 7.4) close to the apparent phase boundary (Figure 3d, left panel). Next, this phenomenon was explored for a sample that forms visible droplets after 1 day (100 μM α-Syn+15% (w/v) PEG-8000). We found that after just 1 round of heating/cooling cycle (within 45 min), the normally delayed LLPS sample phase separated into spherical, liquid-like micron sized assemblies (Figure 3d). However, we could not detect any increase in the SLS intensity upon completing a heating/cooling cycle for most of the nanocluster containing samples (Figure 3d)—likely due to the sensitivity limit of our SLS setup. In order to probe whether temperature-induced size changes of nanoscale clusters can be observed in MP, we studied 10 μM α-Syn+20% (w/v) PEG-8000 solutions at different temperatures by MP. Intriguingly, our data showed that the size of the nanoclusters increased from ~100 kDa to ~400 kDa when the solution temperature was shifted from 25 to 50°C (Figure 3e, left panel). At even higher temperatures (65°C), the nanoclusters shrunk down to ~60 kDa (Figure 3e, right panel)—but did not completely dissolve within the experimental timescale (5 min). Taken together, our observations suggest that the initial nucleation barrier of nanocluster formation is small. However, they reside in a kinetically arrested, local energy minimum which prevents further growth (into micron scale droplets). This kinetic barrier can be overcome by the nanoclusters in either direction (towards monomer or larger assemblies). At higher temperatures (50°C) (or during large incubation times), nanoclusters are enabled to evolve into macroscopic liquid-like assemblies—however, even higher temperatures (65°C) lead to their dissolution (Figure 3e, lower panel).

We wanted to investigate whether the invisible (by optical microscopy) α-Syn nanoclusters accounted for a significant proportion of the volume fraction relative to the dilute phase. One way to probe this was using flow induced dispersion analysis (FIDA)—a technique that can identify small changes in the hydrodynamic radius of protein molecules with great accuracy ^60,82^(Supplementary Figure 7). A plug (few nanoliters) of different α-Syn+PEG-8000 solutions were sent through the FIDA capillary, and the polydispersity index (PDI) was measured from the Gaussian signal profile of individual samples using the in-built FIDA PDI calculator. In FIDA, a PDI value more than 0.05 indicates a polydisperse solution containing heterogeneous assemblies. Our data showed no evidence of polydispersity even for the nanocluster samples (Supplementary Figure 7). More specifically, with increasing α-Syn concentration (in presence of 5% and 20% (w/v) PEG-8000), the PDI values remained almost unchanged and in the order of ~10^-4^(Supplementary Figure 7).

In order to check how the dilute phase concentration of α-Syn changes during the growth of nanoclusters, we centrifuged 50 and 300 μM α-Syn in presence of 20 and 5% (w/v) PEG-8000 (nanocluster forming conditions) at 16000Xg and at different timepoints (day 0-day 7). We observed that the dilute phase concentration remained unaltered even after 7 days of incubation at 25°C (Figure 3f). We reasoned that the nanoclusters might be too small to be pelleted down by normal centrifugation methods routinely used for LLPS^74^ and therefore, used ultracentrifugation^83,84^. Surprisingly, even after ultracentrifuging the samples at 180000Xg, we did not see substantial decrease in the dilute phase concentration with time (Figure 3f). Next, we analyzed whether it is possible to spin down α-Syn nanoclusters under our experimental setup. To this end, we calculated the terminal velocity (V_t_) of a nanocluster by equating the opposing forces (F_c_ = F_b_ + F_f_) during ultracentrifugation (Figure 3g). Our calculations and relevant assumptions are described in detail in the method section of this manuscript and in Figure 4. For a nanocluster with a radius of 30 nm, V_t_ was calculated to be approximately ~2 μm/s. This suggested that at 180000Xg, most of the nanoclusters could indeed be separated from the solution and they would take ~100 minutes time to reach the bottom of the ultracentrifuge tube (1 cm)—which also fits well with our experimental parameters (2h ultracentrifugation). Taken together, our results clearly show that in the nanocluster regime, α-Syn assemblies account for a very small volume fraction at protein concentrations below C_app_, which remains similar throughout the incubation period, and the growth of these assemblies are very slow (scenario ***a***). Intriguingly, in MP, we observed that the nanoclusters could be dissolved upon dilution even after day 2—indicating that they remain liquid-like for a considerable amount of time (scenario ***a***). However, they persisted after dilution after day 4 of incubation (Supplementary Figure 8)—indicating that the nanoclusters gradually solidify just before they appear under the microscope (4-6 days). At this point, we cannot rule out the possibility that after solidification, the nanoclusters become sticky, clump together into bigger assemblies, and reach the resolution limit of conventional microscopy. In this context, we indeed found evidence of clustered α-Syn assemblies in 50 μM α-Syn+20% (w/v) PEG-8000 solution after 4 days of incubation (Supplementary Figure 8).

**Figure 4.**
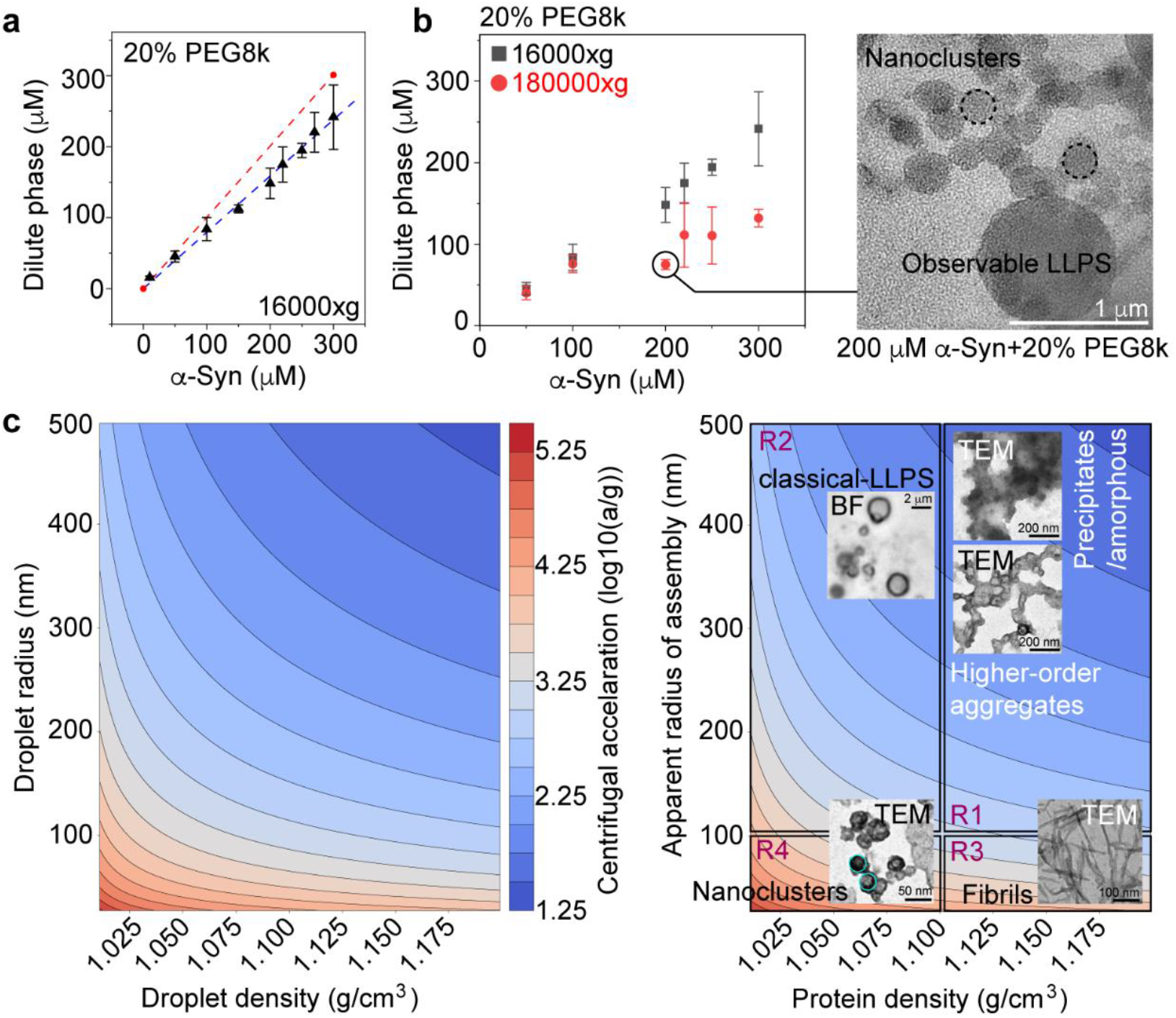
Nanoscale LLPS accounts for a large dense phase fraction above saturation concentration: **a**. Dilute phase concentrations of α-Syn (10-300 μM+20% (w/v) PEG-8000) as a function of the total protein concentration after centrifugation at 16000Xg is shown. The droplets start to appear above ~100 μM total α-Syn concentration. The red dashed line represents the total protein concentration as a guide to the eye. Values represent mean ± S.D. for n=3 independent experiments. **b**. (left panel) Dilute phase concentrations of α-Syn (10-300 μM+20% (w/v) PEG-8000) as a function of the total protein concentration after centrifugation at 16000Xg and 180000Xg is shown. Values represent mean ± S.D. for n=3 independent experiments. (Right panel) TEM images of 200 μM α-Syn+20% (w/v) PEG-8000 showing the presence of both microscopic and nanoscopic droplets above C_app_. The experiment is performed twice with similar observations. **c**. (left panel) A contour plot describing the centrifugal acceleration required to move spherical protein assemblies of different sizes (25-500 nm radius) and different densities (1.01-1.20 g/cm^3^) by 1 cm in 1h in an aqueous solution at 25°C. The color codes represent centrifugal acceleration (RCF) in log (10) scale. (Right panel) The contour plot is divided into 4 regions (R1-R4) to highlight the RCF required to spin down different types of protein assemblies (amorphous aggregates, LLPS droplets, amyloid fibrils and nanoclusters) based on their apparent radius and density. Representative TEM images of α-Syn amorphous and higher order aggregates (R1) (700 μM at pH 5.0), amyloid fibrils (R3) (100 μM, prepared with agitation^85^ at 25°C) and nanoclusters (R4) (50 μM+20% (w/v) PEG-8000) are shown. Representative BF microscopy images of α-Syn LLPS (200 μM+20% (w/v) PEG-8000) is shown for R2. The nanoclusters are highlighted with cyan circles.

### Persistence of nanoscale phase separation above C_app_

Next, we wanted to see whether α-Syn nanoclusters existed even above C_app_ (under LLPS conditions). At 20% (w/v) PEG-8000 concentrations, the C_app_ of α-Syn, i.e., the concentration above which micron sized droplets were visible in the optical microscope, was ~100 μM (Figure 1). Theoretically, for ≥100 μM, the dilute phase concentration should remain constant irrespective of the total protein concentration^34^. We found that unlike expected for a canonical protein system undergoing LLPS, the dilute phase α-Syn concentration, as determined after centrifugation at 16000Xg did not reach a plateau above the critical concentration when the total concentration was increased, which would be expected if the system were to move along a tie line^34^ (Figure 4a). Strikingly, when the samples were ultracentrifuged at 180000Xg, the dilute phase decreased substantially for concentrations above C_app_ and approached a constant value (Figure 4b, left panel). This indicated that even inside the LLPS regime, nanoclusters accounted for a significant volume fraction of the dense phase. TEM imaging of 200 μM α-Syn+20% (w/v) PEG-8000 clearly showed the presence of spherical nanoclusters along with micron-sized droplets (LLPS) in the same sample (Figure 4b, right panel). We also noted that for this LLPS sample, the nanocluster size was substantially larger (radius of ~100 nm)— although remaining marginally below the resolution limit of conventional microscopy to be identified as spherical phase separated assemblies (Figure 4b, right panel).

Our current observations suggest that caution needs to be exercised when using centrifugation to determine the dilute phase concentrations of phase separated samples where nanoscale clusters could be present. In this context, we derived an equation to calculate the centrifugal acceleration (n) required to physically move a spherical protein assembly (of volume V and density ρ_d_) by *h* m within a given time (t) in an aqueous solution (viscosity, *η* = 1 mPa.s). The g-force or RCF value (*n*) can be expressed with the following equation:

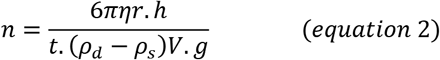

Any protein assembly at sub- or supersaturated concentrations will have a density (ρ_d_) greater than the dilute aqueous phase (ρ_s_≈1.00 g/cm^3^). The upper limit of ρ_d_ is ~1.35 g/cm^3^ for pure protein. Here, we assumed that for micron scale droplets and nanoclusters, the density is likely to be in the range of 1.025-1.100 g/cm^3^. Within this density regime, we define as nanoclusters all objects that cannot be resolved in diffraction-limited microscopy, i.e. tens of nm. On the other hand, higher-order protein assembly/amorphous aggregates (>100 nm) and amyloid fibrils (<100 nm) have densities closer to pure protein (>1.100 g/cm^3^). To obtain equation 2, we took h = 1 cm and t = 1 h and plotted our data in the density (ρ_d_) range of 1.01-1.2 g/cm^3^ (Figure 4c, left panel). We use the term *‘apparent radius of assembly’* (Figure 4c, right panel) which can be defined as the size of a perfect sphere with the same density and sedimentation speed as any of these assemblies. With the help of our analyses, we could reliably obtain four distinct regions with a range of centrifugal acceleration which can be used to isolate amorphous/higher order protein aggregates (R1), classical phase separated droplets (R2), amyloid fibrils (R3) and nanoclusters and nanoscale LLPS (R4) (Figure 4c, right panel). For 20% (w/v) PEG-8000 induced phase separation/cluster formation (such as for α-Syn), the viscosity, *η*, was taken as 20 mPa.s^86^ in equation 2 and plotted (Supplementary Figure 9). Both our observations and calculations indicate that commonly used centrifugation methods (10^3^-10^4^ RCF) are not efficient enough to account for the volume fraction in the nanocluster regime (>10^5^ RCF).

### α-Syn nanoclusters can form in the absence of air-water interfaces

Previously, it has been shown that air-water interfaces and surfaces play a major role in modulating α-Syn assembly ^73,87,88^. To delineate the effect of surfaces, we injected a delayed α-Syn LLPS system (100 μM α-Syn + 15% (w/v) PEG-8000) (Supplementary Figure 1) into a glass capillary and checked for LLPS at different time-points. Our results showed that the first droplets nucleated on the wall of the glass capillary after 24h (Figure 5a, left) and grew substantially larger after 48h (Figure 5a, right) of incubation at 25°C. Intriguingly, just above the C_app_ (100 μM+20% (w/v) PEG-8000), we also observed the appearance of new condensates on the coverslip surface after 4-5h of incubation at 25°C. The delayed, surface nucleated condensates were liquid-like as confirmed by rapid growth/ripening and fusion events upon contact (Figure 5b, Supplementary movie 7). To understand whether delayed cluster/LLPS formation below C_app_ was exclusively due to surface nucleation, we used a 3-component microfluidics-based droplet maker^89,90^ (Figure 5c) to encapsulate α-Syn and PEG without access to a surface with protein affinity. To do this, 600 μM α-Syn and 10% (w/v) PEG-8000 were separately injected into the microfluidic device (yielding a final concentration of 300 μM α-Syn + 5% (w/v) PEG-8000, after mixing) followed by encapsulation of the aqueous reaction mixture by an immiscible fluorocarbon oil (FC-40) into spherical micro-droplets (Figure 5c). The droplets are stabilized by a surfactant that is protein-repellant on the inner droplet surface. Next, these micro-droplets were loaded into a glass capillary and visualized under the microscope at different time-points. We observed that immediately after encapsulation, no microscopic LLPS droplets were present in the protein solution (Figure 5c). Strikingly, like in bulk experiments, we found that distinct, micron sized α-Syn clusters appeared inside the micro-droplets after a week of incubation at 25°C (Figure 5d, Supplementary movie 8). In line with our previous observations (in tubes), the clusters were ThT positive (Figure 5e) and upon further ageing (day 10 and day 15), fibrillar morphology started to appear throughout the micro-droplet (Figure 5e). Notably, 50 μM α-Syn + 20% (w/v) PEG-8000 solution also showed similar results in another independent experiment (Supplementary Figure 10). Taken together, our results clearly suggest that glass surfaces facilitate the nucleation of α-Syn liquid droplets, presumably because of wetting, but that the presence of such surfaces is not a requirement for α-Syn assembly.

**Figure 5.**
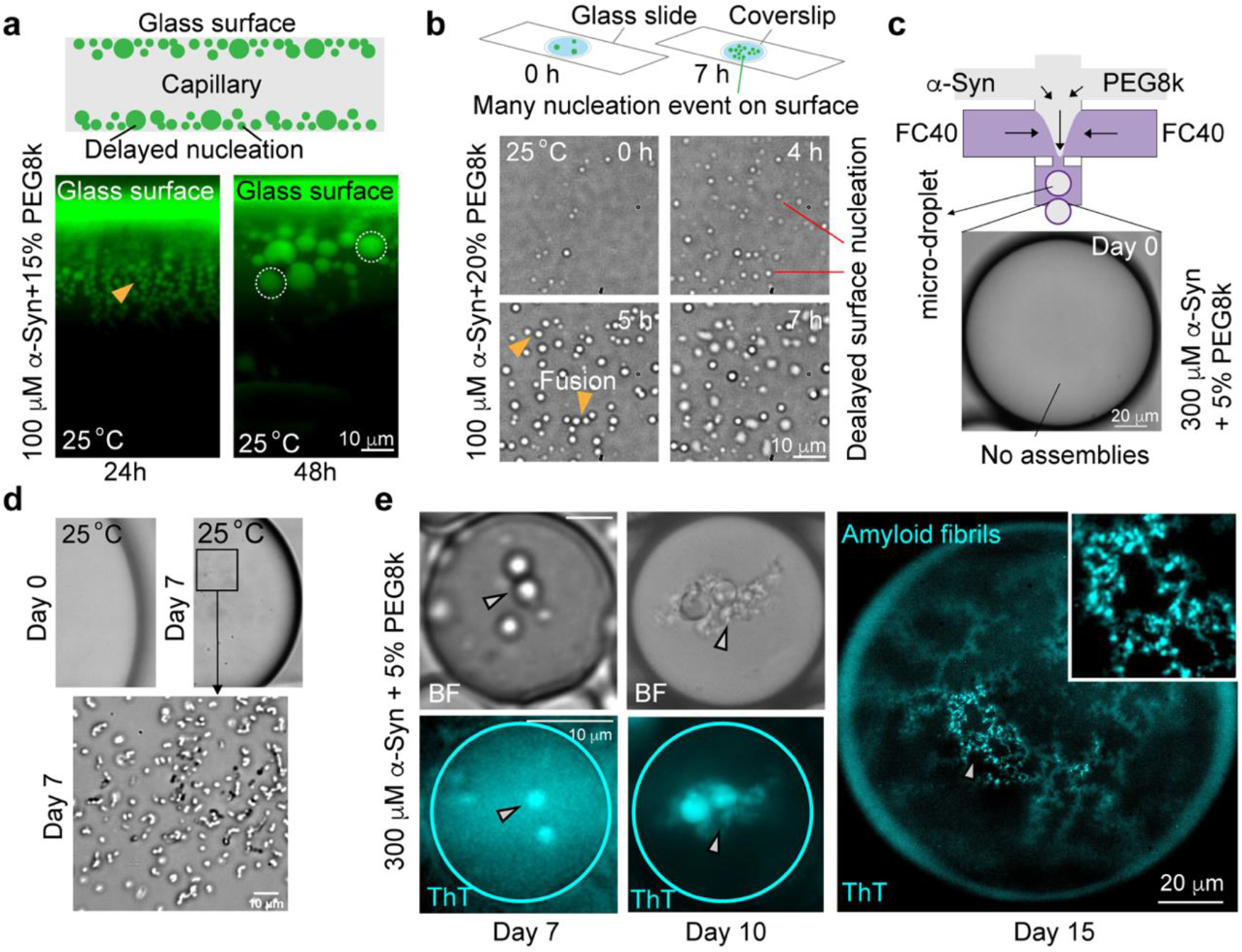
Surfaces can promote delayed α-Syn assembly/LLPS but is not a mandatory requirement: **a**. (upper panel) Schematic showing α-Syn droplets nucleating after a certain period of incubation time on the surface of a glass capillary. (Lower panel) Fluorescence images showing droplet nucleation on the surface of the glass capillary for 100 μM α-Syn + 15% (w/v) PEG-8000 solution (labeled with 100 nM Alexa488-140C-α-Syn) after 1 day (left) and 2 days (right) of incubation at 25°C. The droplets are marked with yellow triangular pointer (day 1) and dashed circles (day 2). The experiment is performed 2 independent times with similar observations. **b**. (upper panel) Schematic showing how new droplets nucleate on glass surface when an already phase separated α-Syn solution is aged for 5-7h on a sealed coverslip. (Lower panel) BF microscopy images of 100 μM α-Syn + 20% (w/v) PEG-8000 solution (sealed in a coverslip) showing delayed nucleation of condensates after ~5h of incubation at 25°C (marked with red lines). Fusion events between the delayed condensates are denoted with yellow triangular pointers. The experiment is performed 2 independent times with similar observations. **c**. (upper panel) Schematic depicting the design of the 3-component microfluidic device. 600 μM α-Syn and 10% (w/v) PEG-8000 are injected in two separate channels. After mixing, the solution is encapsulated with FC-40 which is injected in the third channel. (Lower panel) BF microscopy images of the microdroplets showing no microscopic cluster formation at 0h. **d**. (Upper panels) BF microscopy images showing a microdroplet at day 0 and day 7 (20X magnification). (Lower panel) BF microscopy image of α-Syn clusters that are formed inside the microdroplet after 7 days of incubation at 25°C, shown with a higher magnification (40X). The experiment is performed twice with similar observations. **e**. (left panel) BF and fluorescence microscopy images of the same microdroplet after 7 and 10 days of incubation at 25°C containing 300 μM α-Syn and 5% (w/v) PEG-8000 (spiked with 50 μM ThT). The α-Syn clusters at day 7 and aggregates at day 10 are marked with white triangular pointers. (Right panel) Fluorescence microscopy image of a microdroplet containing 300 μM α-Syn and 5% (w/v) PEG-8000 (spiked with 50 μM ThT) after 15 days of incubation at 25°C. ThT positive, fibrillar morphology has been highlighted as an inset and marked with a white triangular pointer. The experiment is performed twice with similar observations.

## Discussion

The canonical view of LLPS is that if the system is in the binodal zone (metastable), condensate droplets form through a nucleated growth process, whereas they can form through spinodal decomposition once the system is further quenched into conditions where LLPS is more favorable^28,34,37,39^. Inside the spinodal region, LLPS occurs in a barrier-less manner, virtually instantaneously. However, several recent observations do not easily fit into this simplified framework. Notably, it was observed that the rapid (ms) formation of nanoscale assemblies, detectable by stopped flow small angle scattering techniques, precedes the formation of macroscopic droplets, which proceeds on a much slower time scale^37^. Furthermore, it was reported that nanoclusters exist even in regions of the phase diagram outside the droplet formation regions, as defined by microscopically visible droplets^38^. It has been shown that the formation of protein nanoclusters can be, to some extent, decoupled from the formation of macroscopic droplets. There appear to be two energy scales, as well as two-time scales that govern the overall behavior of some proteins prone to undergo LLPS.

In this work, we have been able to study the LLPS behavior of α-Syn in the light of these recent advances and believe that our results make an important contribution to the ongoing discussion. Differently from the other protein systems for which the presence of nanoclusters has been shown (FUS, hnRNPA1)^37,38^, the LLPS of α-Syn displays a very wide range of kinetic time scales, ranging from seconds^59^ to days^58^, depending on the solution conditions (Figure 1). We set out to explore whether the very slow kinetics of demixed assembly formation under some conditions is due to high free energy barriers for droplet nucleation, as would be expected if the system behaves according to classical nucleation theory. In order to probe α-Syn solutions under various conditions for the presence of pre-nucleation clusters and nuclei, we employ a combination of MP, DLS and TEM along with state-of-the-art microscopic techniques in the current body of work.

MP^77^, also known as interferometric scattering microscopy (iSCAT)^78,79^, is a method that has only recently become widely available. It is based on the detection of scattered light that interferes with a reference beam—the incident reflected beam. Although MP measures the mass directly from single molecules (unlike DLS), in simple terms, the scattered intensity depends on the third power of the particle size, compared to the usual sixth power of Rayleigh scattering. This weaker dependence on size renders it possible to detect very small particles, down to individual protein molecules. Indeed, the lower limit in size/molecular weight for individual proteins to be detected is approximately 40 kDa. This cut-off is conveniently placed such that individual α-Syn monomers (14.4 kDa) are not visible, whereas oligomeric assemblies or clusters are expected to be detectable. MP has been successfully used to detect virus particles, protein complexes and protein aggregates^77,91,92^. Here, we employ it, for the first time to our knowledge, to study the phenomenon of LLPS (Figure 2). We prepared solutions of α-Syn under a wide variety of solution conditions, with respect to pH, salt concentration and PEG concentration, under which microscopically visible (i.e., micron sized) droplets and clusters are visible only after a certain time-period (after hours to days). We find that in all scenarios in which we ultimately detect α-Syn droplets and/or assemblies, we can detect nanoscopic assemblies/clusters within the time resolution (seconds) of the MP setup (Figure 2). We have performed a range of control experiments to exclude that the nanoscale clusters we observe originate from processes other than association of α-Syn molecules. Pure PEG or protein solutions do not yield any such clusters. Furthermore, they appear at very low, physiologically relevant protein concentration, even more so, when the pH is titrated down from pH 7.4 towards mildly acidic pH (6.5), despite the titration leading to a slight dilution of the sample.

The dynamic nature of the clusters is demonstrated by dilution experiments, in which they disappear instantaneously. This is in stark contrast to the behavior of more conventional α-Syn oligomers^93,94^, which are highly stable and do not spontaneously disassemble upon dilution but require high denaturant concentrations to dissolve. We therefore describe a new assembly state of α-Syn—nanoscopic, liquid-like clusters (nanoclusters). Very interestingly, recent molecular dynamics simulation studies suggest that α-Syn can form electrostatically driven, micro-phase separated assemblies at physiologically relevant concentrations^95^, below the experimentally probed C_sat_ (reported as C_app_ in this manuscript). Our data, however, suggests that the experimentally observed nanoclusters are electrostatically unfavorable, given the finding that salt is required for their formation. For those samples that contain the smallest observed nanoclusters, we can estimate the number of α-Syn molecules contained in them, by means of a standard calibration of the MP setup with proteins of known MW. This estimate of MW is based on the assumption that the PEG concentration is not significantly different inside the nanoclusters compared to the dilute phase concentration. In this case, the scattering by the nanoclusters stems entirely from their protein content. This assumption may not be always fulfilled, given the recent evidence for direct interactions between proteins undergoing LLPS and PEG^96^. With mass photometric measurements combined with TEM imaging, we find that the α-Syn nanoclusters contain several dozens of molecules with an apparent concentration in the low mM range (Figure 2). However, under some conditions, such as those where we find macroscopic droplets after about 1 day, the clusters display very rapid growth and within seconds grow beyond the upper size limit of the MP setup (5 MDa). Under those conditions, we can follow their growth by conventional solution based DLS. Even though the growth appears very rapid on the nanoscale, it is nevertheless compatible with a time scale of hours to days for the appearance of micron-sized droplets. This can be illustrated by a simple back-of-an-envelope calculation. Let us assume a nanocluster of ca. 100 molecules growing into a droplet of 5 μm diameter. The latter has a protein concentration of several mM, corresponding to somewhere between 10^7^ and 10^8^ molecules of α-Syn. If this size is to be reached within 24h, the average rate of addition of monomers to the cluster is ca. 100-1000 s^-1^. Even though the rate of monomer addition is likely to scale with surface area and therefore expected to increase as the cluster grows, this simple estimate shows that growing within seconds or minutes out of the size range of MP is compatible with a timescale of hours to days to grow towards a micron-sized droplet. The overall behavior of the solution under conditions under which such growth rates are observed correspond closest to the expected behavior of a classically nucleated system. It appears, however, that even under these conditions, and atypical in a classically nucleating system, the kinetics of nucleation is not significantly slower than the growth, as it takes only seconds to form a multitude of clusters within the microscopic field of view. Moreover, SLS and MP measurements at different temperatures suggests that the nanoclusters could reside in a local energy minimum which likely explains their slow growth (Figure 3). The current set of evidence not only shines light on phase separation in the nanoscale regime, but also acknowledges the overwhelming complexity of α-Syn as a phase separating system.

Under nanocluster forming conditions, most conventional experimental methods such as centrifugation followed by measurement of the supernatant concentration and measurement of polydispersity did not reveal their presence (Figure 3). Not only do they require prolonged ultracentrifugation to get spun down, but also, they only correspond to an almost vanishingly small volume fraction of the overall sample (Figure 3). In these cases, it is remarkable that by the time we can observe structures in the optical microscope, the latter are ThT positive, indicating that they contain amyloid fibrils. This observation shows that the kinetics of growth of these particles is of the same order as the formation and growth of amyloid fibrils. The formation of nanoclusters and droplets has been, as mentioned before, associated with two distinct energy and time scales in some protein systems^37^. Our findings suggest that for α-Syn, at least one additional energy and time scale needs to be considered—the formation of amyloid fibrils inside the assemblies irrespective of whether they are nanoscopic or microscopic. To-date, it has mostly been assumed that the kinetics of droplet formation (seconds) is de-coupled from the subsequent formation of amyloid fibrils, because of the much slower timescale of the latter (3-4 orders of magnitude). Here we find that these two processes can happen on the same timescale to an extent that macroscopic liquid droplets are never observed; once they reach microscopically observable sizes they are already largely aggregated. The relative energy scales and timescales of nanocluster formation, droplet formation and amyloid fibril formation can be tuned by the key variables of protein concentration, solution pH, ionic strength and PEG concentration. It is indeed intriguing that under some conditions, the addition of further monomers to a nanoscale cluster seems to have a similar kinetics as the addition to an amyloid fibril. The latter process has been shown to be associated with significant free energy barrier in most cases, which is linked to structural rearrangement and desolvation^97–101^. It remains elusive which physical processes are at play to create a high energy barrier for addition to a nanocluster or small droplet. It is possible that the α-Syn nanoclusters have high interfacial tension and high surface charge^33^ and therefore, it is difficult for new monomers to enter.

The significance of our discoveries discussed in this work stems from the yet unclear role that LLPS plays in pathological protein aggregation. Before the discovery that α-Syn can undergo LLPS and that this process facilitates its amyloid fibril formation, it was thought that α-Syn amyloid fibril formation occurs through heterogeneous nucleation, i.e., it required catalysis or facilitation by suitable surfaces in all cases^71,73^. This includes polymer surfaces, the air-water interface and lipid bilayers. However, it is becoming increasingly clear that amyloid fibrils can form through homogeneous nucleation inside dense condensate droplets. Previous evidence suggested that the nucleation takes place inside micron-scale droplets, but here we find that the nucleation and growth of amyloid fibrils is concomitant with the overall growth of the particles. In order to demonstrate that the nucleation of amyloid fibrils is indeed linked to the formation of the clusters and not catalyzed by surface interactions, we performed a range of additional experiments in the presence and absence of surfaces with a binding affinity for α-Syn. We find that glass surfaces can enhance the formation of droplets and the subsequent amyloid fibril formation (Figure 5). However, we also perform experiments inside water-in-oil emulsion droplets that are stabilized by a protein repellent surfactant. Under conditions where we observe nanoclusters, we also observe the emergence of ThT positive α-Syn assemblies over time which agrees with the time scale of their appearance in bulk solution (Figure 5).

Taken together, we have discovered nanoscopic clusters of α-Syn by MP that can act as precursors to both macroscopic droplet formation, as well as amyloid fibril formation. This insight is valuable in the context of the open questions as to the link between protein LLPS and amyloid fibril formation. Furthermore, we believe the features of MP to be uniquely suitable for the study and quantification of nanoscopic LLPS precursors and our work therefore to be of wide interest to the community concerned with protein LLPS.

## Materials and Methods

### Materials

pT7-7 plasmid encoding the human α-synuclein (α-Syn) (*SNCA*) was sequenced by GATC (Eurofins, Germany) to ensure no point mutations had occurred. LB broth and ampicillin for wild type (WT) α-Syn purification were purchased from VWR (Denmark). Purified Alexa-488 maleimide labeled 140C-α-Syn (24 μM stock solutions prepared in 10 mM sodium phosphate buffer (NaH_2_PO_4_.H_2_0+Na_2_HPO_4_.2H_2_0) pH 7.4) was a kind gift from Celine Galvagnion Buell (University of Copenhagen). NaH_2_PO_4_.H_2_0, Na_2_HPO_4_.2H_2_0, PBS tablets, NaCl, PEG-8000 and PEG-3000 was procured from Sigma (USA). Ficoll-400 was purchased from ThermoFisher scientific (USA). The buffer compositions, NaCl, PEG and Ficoll were dissolved in RNAse and nuclease free milliQ water. Thioflavin-T (ThT) was purchased from Sigma (USA). Formvar/carbon coated TEM grids were procured from Sigma (USA). All other relevant reagents/materials were purchased from Sigma (USA) and VWR (Denmark) unless otherwise specified.

### Methods

#### Expression and purification of recombinant Wild-type (WT) α-Syn

5 ml of primary of *E. coli* BL21 (DE3) culture (grown for 10-12h at 37°C) with the pT7-7 plasmid carrying WT human α-Syn gene (*SNCA*) was inoculated into 1 L of LB with 100 mg/L ampicillin (Amp) as the selection marker. The culture was grown in a 3 L flask at 37°C with a shaking speed of 180 rpm. The OD_600_ was measured at regular time intervals till it reached 0.8 and protein expression was induced by addition of 1 mM isopropyl ß-D-1 thiogalactopyranoside (IPTG). After IPTG addition, the cells were incubated at 37°C with 180 rpm shaking for 4h and subsequently harvested by centrifugation (7000Xg, 20 min, 4°C). The cell pellets obtained from 1 L culture were stored at −20°C until further use. For purification, cell pellet from 1 L culture was dissolved in 20 ml of 10 mM Tris-HCl, 1mM ethylenediamine tetra acetic acid (EDTA), pH 8.0 with 1mM phenylmethylsulfonyl fluoride (PMSF) and subsequently sonicated on ice for 2 min (10 s sonication, 30 s pause, 12 rounds at 40% amplitude). Immediately after sonication, 1μL Benzonase was added to the cell lysate to precipitate DNA. The solution was then centrifuged at 20000Xg for 30 min at 4°C and the supernatant was collected and boiled on a water-bath for 20 min. After boiling, the solution was cooled down at room temperature and centrifuged at 20000Xg for 20 min at 4°C. This step was performed in order to precipitate heat-sensitive proteins while α-Syn remained in the supernatant. The supernatant was collected, the volume measured and 4 ml saturated (NH_4_)_2_SO_4_ was added for 1 ml supernatant to salt out α-Syn. The salted out α-Syn was isolated by centrifuging at 20000Xg for 20 min at 4°C. The supernatant was discarded, and the protein pellet was dissolved in 7 ml of 25 mM Tris-HCl, pH 7.7. 7μl dithiothreitol (DTT) was added to a final concertation of 1 mM to the solution. Subsequently, the α-Syn solution was dialyzed against 25 mM Tris-HCl, pH 7.7, 1 mM DTT (the same buffer) for 18h at 4°C to remove excess (NH_4_)_2_SO_4_. During dialysis, the tank was replenished with fresh buffer after 12h. Next, the α-Syn solution was subjected to anion exchange column (AEC) (HiTrap Q Hp 5 ml, GE healthcare) followed by size exclusion chromatography (SEC) (HiLoad 16/600 Superdex 200 pg. column). Finally, the protein was eluted in 10 mM of sodium phosphate (NaH_2_PO_4_.H_2_0+Na_2_HPO_4_.2H_2_0) buffer (pH 7.4). α-Syn concentrations in the final elution was assessed using a UV spectrophotometer—ProbeDrum (ProbationLabs, Sweden) by measuring the absorption at 280 nm. The theoretical molar extinction coefficient of WT α-Syn (5960 M^-1^cm^-1^) was predicted by ProtParam (Expasy, Switzerland).

#### In vitro liquid-liquid phase separation (LLPS) of α-Syn

The AEC/SEC purified stock α-Syn solutions were concentrated in 20 mM PBS (20 mM NaH_2_PO_4_.H_2_0, Na_2_HPO_4_.2H_2_0, 250 mM NaCl, 5 mM KCl, pH 7.4) using 3 kDa filters (500 μl Amicon ultra centrifugal cutoff filters) unless mentioned otherwise. Samples containing different concentrations of α-Syn (10-300 μM) and PEG-8000 (5-20% (w/v)) were prepared in 20 mM PBS, pH 7.4, at room temperature (25°C) in 1.5 ml Eppendorf tubes. From these samples, 5 μl of each solution was drop casted on pre-cleaned 76 mm x 26 mm x 1 mm glass slides (Epredia, USA) and imaged under a microscope. The technical details of the microscopes are provided in the subsequent section(s). For fluorescence and/or confocal microscopy, individual samples were spiked with 100 nM Alexa-488-140C-α-Syn and/or 50 μM Thioflavin-T (ThT) before addition of PEG-8000. For time-lapse movies (experiments described in Figure 1e and Figure 5b), 5 μl of the phase separated solution was drop casted on a clean glass slide and sandwiched with a 18 mm glass coverslip. The edges of the coverslip was subsequently sealed using commercially available capillary sealing paste (NanoTemper Technologies, Germany) to prevent evaporation. The samples were then subjected to time-lapse imaging under bright field (BF) or fluorescence microscopy at 25°C. For the experiment described in Figure 5a, 100 μM α-Syn + 15% (w/v) PEG-8000 spiked with 100 nM Alexa-488-140C-α-Syn was prepared in 20 mM PBS (pH 7.4) and loaded into a hollow square glass capillary (0.70 mm x 0.70 mm) (CM Scientific, Ireland) and imaged at different timepoints (day 0-2) by a epifluorescence microscope at 25°C. To note, the sample was incubated in the glass capillary by sealing the ends with the help of a commercially available sealing paste (NanoTemper Technologies, Germany). Similar protocol was exercised to visualize aggregation in 700 μM α-Syn (prepared in 20 mM PBS (pH 7.4) under a glass coverslip after 7 days incubation at 25°C. The appearance of aggregates within 3-5h of incubation in this sample was likely due to presence of the glass surface promoting clumping of the fibrils (Supplementary Figure 1). To check delayed cluster formation below the apparent critical concentration (C_app_), 300 μM α-Syn + 5% (w/v) PEG-8000 and 50 μM α-Syn + 20% (w/v) PEG-8000 samples were prepared in 20 mM PBS (pH 7.4) and incubated in 1.5 ml Eppendorf tubes for 7 days at 25°C. The tubes were wrapped with parafilm to prevent evaporation during the incubation period. At different timepoints (day 0-7), 5 μl of the solution was drop casted onto clean glass slides and imaged under a microscope. In a parallel experiment, identical samples were spiked with 100 nM Alexa-488-140C-α-Syn and/or 50 μM ThT and observed after 7 days of incubation at 25°C with the help of a fluorescence microscope. To note, 0.05% sodium azide was added to all samples to prevent bacterial/fungal contamination over long incubation periods.

#### Bright field microscopy

Bright field (BF) microscopy images reported in this manuscript were captured using an inverted bright field microscope (Zeiss Axio vert. A1, (Zeiss, Germany)) with a 40X objective lens unless mentioned otherwise. For time-lapse BF microscopy experiments (Figure 5a), images from 100 μM α-Syn + 20% (w/v) PEG-8000 solution (prepared in 20 mM PBS, pH 7.4) were acquired every 2 minutes for ~30h at 25°C. Time-lapse BF movies of the α-Syn clusters formed in 300 μM α-Syn + 5% (w/v) PEG-8000 and 50 μM α-Syn + 20% (w/v) PEG-8000 samples were captured with an interval of 800 ms at 25°C unless mentioned otherwise. All BF microscopy images were obtained at 16-bit depth with a resolution of 1024 x 1024. The images were subsequently analyzed with ImageJ (NIH, USA).

#### Fluorescence microscopy

ThT Fluorescence images of the α-Syn droplets/clusters was imaged using an in-built ThT channel (ex. 436/20, beam splitter 455, emission 480/40) in the Zeiss Axio vert. A1 (Zeiss, Germany) microscope with a 40X objective lens unless mentioned otherwise. For the experiment described in Figure 1e, 200 μM α-Syn + 20% (w/v) PEG-8000 solution (LLPS) spiked with 50 μM ThT was imaged on the glass slide for ~30h at 25°C with an acquisition interval of 2 min. The exposure was set at 800 ms in the ThT channel. All ThT fluorescence images were taken in 16-bit depth with a resolution of 1024 × 1024 and subsequently analyzed using ImageJ (NIH, USA). Identical technical parameters were used for experiments described in Figure 5e and Supplementary Figure 10. Fluorescence images of the α-Syn droplets/cluster samples spiked with 100 nM Alexa-488-140C-α-Syn were captured by an epifluorescence microscope (Nikon Eclipse Ti2 (RAMCON, Denmark)) with the help of 40X and 60X (oil immersion) magnification objectives. The excitation wavelength was set at 470 nm and images were captured with an emission channel in the range of 500-600 nm. The laser exposure was adjusted for individual sample so that maximum number of droplets/clusters can be detected. Droplet fusion (Figure 1b) was captured in 200 μM α-Syn + 20% (w/v) PEG-8000 solution (0h) (spiked with 100 nM Alexa-488-140C-α-Syn) with an acquisition interval of 1 s and with an exposure of 50 ms. Delayed LLPS in 100 μM α-Syn + 15% (w/v) PEG-8000 (in 20 mM PBS, pH 7.4) (spiked with 100 nM Alexa-488-140C-α-Syn) was observed by loading the sample into a hollow square glass capillary (0.70 mm x 0.70 mm) (CM Scientific, Ireland) and imaged at different timepoints (day 0-2) at 25°C at 100 ms exposure with a 40X objective lens (Nikon Eclipse Ti2 (RAMCON, Denmark)). All images were obtained with a 16-bit depth and at a resolution of 2048 x 2048 at room temperature (25°C). Fluorescence imaging was performed at Denmark Technical University (DTU) bio-imaging core facility. The images were subsequently processed and analyzed using ImageJ (NIH, USA).

#### Confocal fluorescence microscopy and photobleaching experiments

An inverted confocal fluorescence microscope (LMI-005-Leica Microsystems Confocal Microscope SP8) was used for the fluorescence recovery after photobleaching (FRAP) experiments described in Figure 1 and Supplementary Figure 1. The samples were imaged with the help of a 60X (oil immersion) objective. For FRAP, a bleaching radius of 1-2 μm was chosen depending on the α-Syn droplet/cluster size. The droplet/cluster was photobleached with a 488 nm laser at 100% power for 2 s. The fluorescence recovery was recorded for 30-90 s (post-bleach) and subsequently corrected for the effect of passive bleaching and the background fluorescence following previously established protocols^58^. The fluorescence recovery (post-bleach) was normalized with respect to the pre-bleach fluorescence intensity for individual α-Syn droplet/cluster^58,60^. All FRAP experiments were carried out at room temperature (25°C). FRAP experiments were performed at DTU bio-imaging core facility. The images were subsequently processed and analyzed using ImageJ (NIH, USA).

#### Transmission electron microscopy (TEM)

TEM imaging of α-Syn phase separated assemblies, nanoclusters, fibrils and amorphous aggregates/precipitates was performed following previously established protocols^58,59^ with slight modifications. Briefly, 10 μl of α-Syn solution under different conditions was drop casted onto a formvar/carbon coated TEM grid (Sigma, USA) and kept standstill for 10 min. The excess solution was then blotted carefully using a Whatman filter paper without touching the grid. Next, 10 μl of 1% (w/v) uranyl acetate solution was drop casted on the grid and the samples were stained for 20-30 s. The excess stain is blotted, and the grids were air dried for 10 min at 25°C before imaging. The grids containing different α-Syn assemblies were subsequently imaged using a 200 kV Tecnai T20 G2 electron microscope (FEI, USA) and at desired magnifications (4000-10000X). The images were captured using a TVIPS XF415 CMOS 4K camera and TVIPS EMplify v0.4.5software (50-100 ms exposure time depending on the sample). TEM imaging was performed in DTU nanolabs imaging facility.

#### Bulk ThT aggregation kinetics of α-Syn LLPS sample

200 μM α-Syn + 20% (w/v) PEG-8000 solution (prepared in 20 mM PBS, pH 7.4, 0.05% sodium azide) was spiked with 50 μM ThT and 100 μl of the solution was loaded into a clear-bottom 96 well plate (Fisher scientific). Amyloid aggregation was monitored at 25°C in a fluorescence microplate reader—FLUOstar Omega (BMG Labtech, Germany) by exciting the sample at 440 nm and recording the emission at 480 nm, at regular intervals (every 15 min). The experiment was performed under quiescent (without shaking) conditions. The total duration of this experiment was ~3 days (Figure 1).

#### Mass photometric measurements and analysis of α-Syn assembly

All mass photometry measurements were carried out using the Refeyn One^MP^ mass photometer (Refeyn Ltd., UK). Rectangular, high precision glass coverslips (24 mm X 50 mm) (Marienfeld) were cleaned following the procedure described by Refeyn Individual rinsing protocol. The coverslips were subsequently placed on the 100X oil-immersion objective and a self-adhesive silicone culture well (Sigma Aldrich GBL103250, Grace Bio-Lab reusable CultureWell gaskets) with 4 wells is placed on top of the coverslip to load the samples. 10-20 μl total volume can be loaded in each well. 2 light sources were used in MP—the primary illumination laser of 525 nm (1 W) and an autofocus laser of 635 nm (3.5 mW). For our experiments (Figure 2, Supplementary Figure 2), 10 μl of α-Syn solutions (without PEG) in 20 mM PBS (pH 7.4, unless mentioned otherwise) were first drop casted onto individual wells and the focal plane was adjusted using the autofocus laser. This autofocusing step is crucial since MP mass measurements are performed based on individual binding events on the coverslip plane (protein(s) binding to the glass surface)^77^. Next, desired volume (in μl) of PEG-8000, PEG-3000 and Ficoll-400 (F-400) solutions prepared in the same buffer (20 mM PBS, pH 7.4) were added at appropriate concentrations. Immediately after addition of the crowding agent(s), mass measurements were carried out using the primary laser for 60 s. 6000 frames were acquired under optimized auto-exposure settings and with the ‘large’ camera acquisition mode with a binned pixel dimension of 128 x 128. The dimension of the entire frame was 10.8 μm x 10.8 μm. The effective detection area was slightly less (95 μm^2^), given the frame-dimensions. To note, the standard curve calibration was performed in a similar manner (Supplementary Figure 2). However, unlike α-Syn (14.4 kDa) and/or PEG (3 or 8 kDa), which were below the detection limit of MP (lower detection limit ~40 kDa), the standard protein samples could be detected (molecular weight (MW) range of ~70-700 kDa). Therefore, initial focusing was performed using 10 μl buffer solution followed by addition of desired volumes of standard protein samples to a final concentration of 50 nM. For the pH titration experiment (Supplementary Figure 2), the focal plane was adjusted with 10 μl of 20 μM α-Syn+10% (w/v) PEG-8000 (pH 7.4). The same sample was measured in a stepwise manner after each 0.5 μl addition of 50 mM HCl. For aged α-Syn nanoclusters (Figure 3) and temperature dependent experiments, the initial focusing was carried out with corresponding PEG-only solutions at identical PEG concentrations as in the samples. The PEG-solutions were carefully pipetted out ensuring minimal disturbance, washed with buffer and nanocluster solutions (differently aged or incubated at different temperatures) were drop casted and immediately recorded. All measurements were performed at 25°C unless mentioned otherwise (e.g., temperature screening experiments). All datasets were recorded with the Acquire^MP^ (v2.4.0) software.

The aim of the initial autofocusing method in MP is to remove the background of the coverslip surface (removal of static background^77^)—thereby allowing quantification of any α-Syn assembly as dark spots/contrast which is termed as ‘ratiometric’contrast^77^. The extent/darkness of the ratiometric contrast can be converted to MW from the standard curve (known ratiometric contrast for known MW) using the Discover^MP^ (v2.304) analysis software. Interestingly, for α-Syn nanoclusters, due to their constant attachment and detachment on the surface, we could visualize individual assemblies for prolonged timescales. The events (spots) captured in each frame gradually populated the contrast/MW scale producing a histogram, which was fit to a Gaussian to calculate the average MW of the assemblies using Discover^MP^ (v2.304) analysis software. Samples closer to the canonical LLPS regime often showed a tailing in the MW histogram likely due to presence of larger assemblies beyond the analytical limit of MP—which was not taken into account for reporting the apparent MW of the nanoclusters.

#### Dynamic Light Scattering (DLS)

Measurements were carried out using a high sensitivity and high throughput capillary based DLS instrument—Prometheus PANTA (NanoTemper Technologies, Germany). α-Syn solutions (monomer, nanoclusters, LLPS) were prepared in 1.5 ml Eppendorf tubes and loaded into 10 μl high sensitivity microcapillaries (NanoTemper Technologies, Germany) and subjected to DLS measurements. The DLS laser (630 nm) was set to 100% power and all size analysis measurements were performed at 25°C. Ten individual iterations were performed for one sample per experiment. The autocorrelation functions from individual samples were used to obtain the size distribution profile by an in-built algorithm. For analysis of the α-Syn assemblies during time (Supplementary Figure 4), individual samples were incubated in 1.5 ml Eppendorf tubes (at 25°C) and 10 μl of the sample was loaded into the DLS capillaries just before measurements at indicated timepoints (day 0-7). The monomer peak was integrated in the range of 1-10 nm in the experiments reported in Supplementary Figure 4 for 50 μM α-Syn+5% (w/v) PEG-8000 and 300 μM α-Syn+5% (w/v) PEG-8000 (20 mM PBS, pH 7.4). To note, since α-Syn nanoclusters were marginally below the detection limit of DLS and 20% (w/v) PEG-8000 remained invisible (Supplementary Figure 3), no viscosity or other relevant correction was necessary for a qualitative analysis of our datasets as described in this manuscript.

#### Static light scattering (SLS) experiments

SLS experiments were carried out using a thermally controlled, multichannel spectrophotometer—ProbeDrum (ProbationLabs, Sweden). A total volume of 70 μl for each sample was scanned in the range of 12-55°C using a 10×4 mm light path quartz glass cuvette (Hellma Analytics, Germany). Measurements were carried out under 3 independent channels: **1**. Absorbance at 240-700 nm. **2**. Fluorescence at 240-700 nm (excitation at 280 nm) and **3**. SLS at 90° with a 636 nm laser. Temperature scan was performed with a 20 s equilibration time and 20 s interval time between each °C. All data were analyzed with the help of PDviewer software (ProbationLabs, Sweden) and plotted using Origin Pro 2019 (Origin labs, USA).

#### Polydispersity index (PDI) measurements using flow induced dispersion analysis (FIDA)

PDI values were obtained from Taylor grams^82^ obtained from running 20-300 μM α-Syn in presence of 20% and 5% (w/v) PEG-8000 (in 20 mM PBS, pH 7.4) at 25°C. Each sample was spiked with equal amount (50 nM) of Alexa-488-140C-α-Syn as a fluorescent reporter. The FIDA parameters used for our experiments are described in the following tables (Supplementary Table 1–2).

**Supplementary Table 1.** FIDA analysis for PDI determination of different concentrations of α-Syn in presence of 20% (w/v) PEG-8000. Tray 1, 2 and the capillary were maintained at 25°C for all our measurements.

| Tray | Vial | Pressure<br>(mbar) | Time (s) | Outlet | Measure | Comment |
| --- | --- | --- | --- | --- | --- | --- |
| 2 | 1 | 3500 | 180 | Variable | no | 1 M NaOH wash |
| 2 | 2 | 3500 | 120 | Variable | no | MQ water wash |
| 2 | 3 | 3500 | 60 | Variable | no | 20 mM PBS buffer wash |
| 1 | Analyte (position 1) | 3000 | 60 | Variable | no | 20 mM PBS with 20% (w/v) PEG-8000 |
| 1 | Indicator (variable) | 1200 | 10 | Variable | no | 20-50 µM α-Syn in presence of 20% (w/v) PEG-8000.<br>Each sample was spiked with 50 nM reporter molecule. |
| 1 | Analyte (position 1) | 3000 | 500 | Variable | yes | 20 mM PBS with 20% (w/v) PEG-8000 |

**Supplementary Table 2.** FIDA analysis for PDI determination of different concentrations of α-Syn in presence of 5% (w/v) PEG-8000. Tray 1, 2 and the capillary were maintained at 25°C for all our measurements.

| Tray | Vial | Pressure<br>(mbar) | Time (s) | Outlet | Measure | Comment |
| --- | --- | --- | --- | --- | --- | --- |
| 2 | 1 | 3500 | 180 | Variable | no | 1 M NaOH wash |
| 2 | 2 | 3500 | 120 | Variable | no | MQ water wash |
| 2 | 3 | 3500 | 60 | Variable | no | 20 mM PBS buffer wash |
| 1 | Analyte (position 1) | 700 | 120 | Variable | no | 20 mM PBS with 5% (w/v) PEG-8000 |
| 1 | Indicator (variable) | 600 | 10 | Variable | no | 20-300 μM α-Syn in presence of 5% (w/v) PEG-8000.<br>Each sample was spiked with 50 nM reporter molecule. |
| 1 | Analyte (position 1) | 700 | 500 | Variable | yes | 20 mM PBS with 5% (w/v) PEG-8000 |

The PDI values were determined using the PDI calculator in the FIDA software suite (v4.3) (FIDA biosystems Aps, Denmark) from corresponding apparent hydrodynamic radius ( $R_h$ ) and $t_R$ (arrival time of the protein peak at the detector)<sup>82</sup>.

#### Quantification of the dilute phase using (ultra)centrifugation

Ultracentrifugation of α-Syn nanoclusters and LLPS samples were performed using an Optima series (Beckman Coulter, USA) ultracentrifuge. 50 μl of each sample was centrifuged at a speed of 180000Xg (180000 RCF) in 7 mm x 20 mm polycarbonate centrifugation tubes (Beckman Coulter, USA) for at least 2h. It is important to note that for 300 μM α-Syn + 5% (w/v) PEG-8000 and 50 μM α-Syn + 20% (w/v) PEG-8000 (in 20 mM PBS, pH 7.4) samples, the conditions in the supernatant remained conducive for α-Syn nanocluster formation even after ultracentrifugation. Reappearance of higher order α-Syn assemblies in the supernatant could result in errors in concentration measurements. Therefore, immediately after centrifugation, 5 μl of the supernatant was diluted below the nanocluster regime (as observed in our MP experiments described in Figure 2) with appropriate dilution factors. The concentration of the diluted supernatant was measured using a NanoDrop Lite (ThermoFisher scientific, USA) and up scaled (multiplied) by its respective dilution factor. This operation was performed for all samples irrespective of whether they were below or above the C_app_ of α-Syn for a given PEG-8000 concentration (at 25°C).

To check whether α-Syn nanoclusters can be spun down using ultracentrifugation at 180000Xg, we assumed that the density of the nanocluster (ρ_d_) = 1.100 g/cm^3^ (at 25°C) and the displacement length (*h*) to reach the bottom of the centrifuge tube = 1 cm. The density of the solvent (20% (w/v) PEG-8000) (ρ_s_) was taken as = 1.01 g/cm^3^ (at 25°C)^86^. Now, the centrifugal force (F_c_) acting on a nanocluster having a radius (*r*) = 30 nm (from TEM imaging, Figure 2) will be given by the following equation (equation 3):

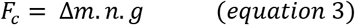

Where the term (n.g) is the RCF centrifugation acceleration (in our case, 180000 times g = 180000 × 9.81 m/s^2^) and Δm is the apparent mass of the protein-rich nanocluster after buoyancy corrections. Therefore, Δm can be described as (equation 4):

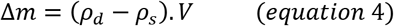

Where ρ_d_ is the density of protein inside the nanocluster; ρ_s_ is the density of the solvent and V is the volume of the nanocluster. The volume of the nanocluster was calculated to be V = 1.13e-16 cm^3^. From there, the F_c_ was quantified as F_c_ = 1.8e-11 g.m/s^2^.

The opposing force that will delay the centrifugation of the nanocluster is the drag or frictional force (F_f_), which can be described by the following equation (equation 5):

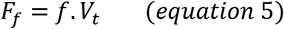

Where, V_t_ is the terminal velocity and *f* is the frictional coefficient (equation 6):

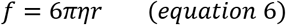

Where r = 30 nm and η= 20 mPa.s = 20 g/m. s (for 20% (w/v) PEG-8000)^86^. The value of *f* was calculated to be = 1.2e-5 g/s.

By equating the two opposing forces, we get: 1.8e-11 = 1.2e-5 (V_t_).

Therefore, the terminal velocity at 180000Xg is = 1.5e-6 m/s = 1.5 μm/s ~ 2 μm/s.

Thus, to travel 1 cm towards the bottom of the tube, the nanodroplet will take ~100 minutes time which is also what we used in our experiment (2 h).

We further extended our interpretation to develop an equation to calculate the RCF centrifugal acceleration (n) required to physically move a spherical protein assembly (of volume, V) by *h* m within a given time (t) in an aqueous solution (viscosity, *η*) (Figure 4 and Supplementary Figure 9). The RCF value (*n*) can be expressed with the following equation (equation 7):

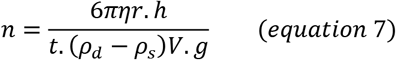

The data was plotted using a python script running in matplotlib^102^ v3.5.0 which can be found with the code availability statement. To note, for all other dilute phase concentration related experiments (Figure 3 – 4), normal centrifugation was carried out for 300 μM α-Syn + 5% (w/v) PEG-8000 and 50 μM α-Syn + 20% (w/v) PEG-8000 (in 20 mM PBS, pH 7.4) samples using an Eppendorf 5418R centrifuge at 25°C.

#### Microdroplet encapsulation to delineate the effect of surfaces on α-Syn LLPS/nanocluster formation

Fluorocarbon Oil-40 (FC-40) encapsulated microdroplets containing α-Syn +PEG-8000 solutions were produced by flow-induced breakup in a microfluidic channel^89,90^. Devices were fabricated according to conventional methods from poly-(dimethyl siloxane)^90^ and immediately following bonding of the device to the glass substrate, a solution of 1H,1H,2H,2H-perfluorododecyl trichlorosilane (0.1 g/L, 0.15 mM) in Fluo-Oil 40 (FC-40) (Emulseo, France) was injected into the device by hand. Following incubation at 80°C for 15 min, the excess reagent was flushed out with FC-40 and the devices were used immediately. Solutions of desired concentration of α-Syn (spiked with 50 μM ThT) and PEG-8000 in an aqueous phase (20 mM PBS, pH 7.4) were separately injected into the central channel under a pressure of 200 mbar (Elveflow OB1 microfluidic flow controller, Elveflow, France). The FC-40 was injected in parallel, with a pressure of 400 mbar. The microdroplets were collected at the outlet and injected into a hollow square glass capillary (0.70 mm x 0.70 mm) (CM Scientific, Ireland) for subsequent imaging (BF and ThT channel) under the microscope (Zeiss Axio vert A1, Zeiss, Germany) equipped with a CFP filter cube (model no. 424931, ex. 436/20, beam splitter 455, emission 480/40), illuminated using a Visitron Cool LED pE100 (Visitron Systems, Germany) operating at 440 nm.

### Amino acid sequences

#### Human α-Synuclein [*SVC4*]

MDVFMKGLSKAKEGVVAAAEKTKQGVAEAAGKTKEGVLYVGSKTKEGVVHGVATVAEKTKEQVTNVGGAVVTGVTAVAQKTVEGAGSIAAATGFVKKDQLGKNEEGAPQEGILEDMPVDPDNEAYEMPSEEGYQDY EPEA

#### A140C α-Synuclein

MDVFMKGLSKAKEGVVAAAEKTKQGVAEAAGKTKEGVLYVGSKTKEGVVHGVATVAEKTKEQVTNVGGAVVTGVTAVAQKTVEGAGSIAAATGFVKKDQLGKNEEGAPQEGILEDMPVDPDNEAYEMPSEEGYQDY EPEC

## Data availability

The authors declare that all the data supporting the findings of this study are available within the paper and in supplementary information files. All the data analysis was performed using published tools and packages and has been cited in the paper and supplementary information text. No data has been excluded.

## Code availability

RCF centrifugal acceleration related analysis was carried out using a python script in matplotlib v3.5.0. The code is provided in the following section:

### Code

~~~
import numpy as np
import matplotlib.pyplot as plt
def n(rho_p, r_p):
    V_p =3.1415*4/3*r_p**3*1e6
    x = 0.01
    t = 3600
    g = 9.82
    rho_s = 1
    eta = 1
    f = 6*3.1415*r_p*eta
    return x*f / (t*(rho_p-rho_s)*V_p*g)
radii = np.linspace(25e-9, 500e-9, 100)
rho_ps = np.linspace(1.01, 1.2, 100)
ns = [n(rho_p, r) for r in radii for rho_p in rho_ps]
fig, ax = plt.subplots()
c =ax.contourf(np.log10(np.reshape(ns,(100,100))), cmap=“coolwarm”, levels = 15,
               extent =[min(rho_ps), max(rho_ps), min(radii), max(radii) ],
               origin=“lower”)
plt.xlabel (“Droplet density [g/cm^3]”)
plt.ylabel (“Droplet radius [m]”)
cbar = plt.colorbar(c) cbar.set_label(“log10(a/g)”) plt.yscale(“log”)
plt.savefig(“img.svg”)
plt.show()
~~~

## Acknowledgements

We thank Prof. Celine Galvagnion Buell (Department of drug design and pharmacology) at Copenhagen University for donating the Alexa488-140C-α-Syn protein. DTU bio-imaging core at DTU bioengineering is acknowledged for confocal/fluorescence imaging and FRAP experiments. DTU nanolabs is acknowledged for TEM imaging. We thank Louise K. Klausen and Kristina Mielec for technical assistance in α-Syn expression and purification. S.R. and A.K.B. would like to acknowledge VILLUM FONDEN for financial support (Grant number 35823). T.O.M. and A.K.B. would like to acknowledge the Novo Nordisk Foundation for funding (Grant number: NNFSA170028392).

## Author contributions

S.R., T.O.M., L.B.T. designed and performed experiments and analyzed data. N.J. analyzed data. A.K.B. acquired funding, conceived and supervised the study, designed experiments and analyzed data. S.R. and A.K.B. wrote the manuscript. S.R. prepared all schematics and illustrations. All authors commented on the manuscript.

