## Supplementary information for "Mass photometric detection and quantification of nanoscale α-synuclein phase separation"

### Supplementary Figures

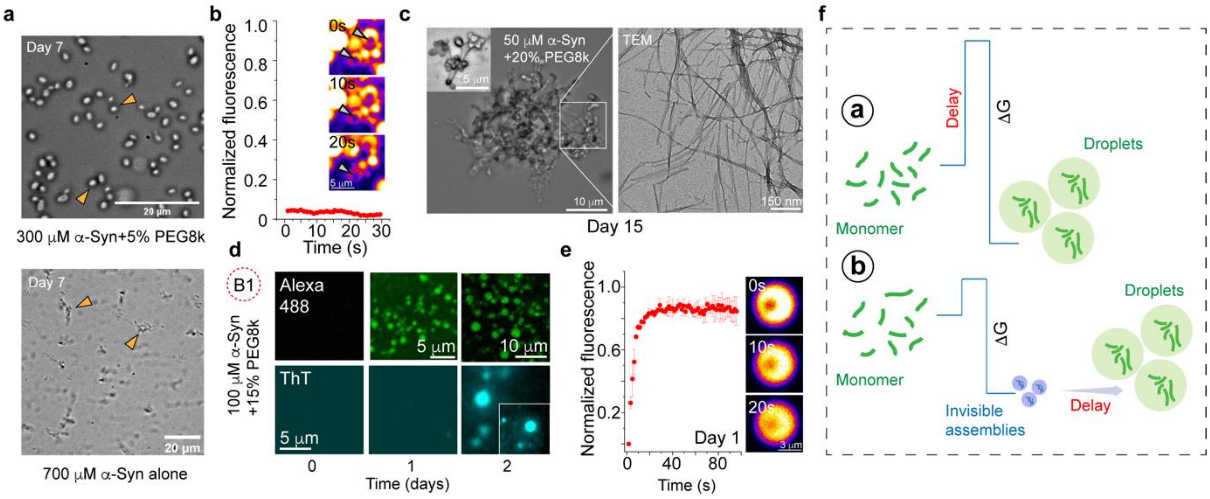


**Supplementary Figure 1. Delayed clustering/LLPS of α-Syn below $C_{app}$: a.** (upper panel) Representative bright field (BF) microscopy image of 300 µM α-Syn in the presence of 5% (w/v) PEG-8000 after 7 days of incubation at 25ºC. (Lower panel) Representative BF microscopy image of 700 µM α-Syn after 7 days of incubation at 25ºC. The α-Syn clusters and aggregates are shown with yellow triangular pointers. The experiments are performed 2 times with similar observations. **b.** Normalized fluorescence recovery after photobleaching (FRAP) of microscopic α-Syn clusters just after their appearance is shown. (Inset) Representative snapshots of a α-Syn cluster at different times post-bleaching. The images are shown in thermal LUT for better visualization. The experiment is performed 2 independent times with similar results. **c.** (left panel) Representative BF microscopy images of 50 µM α-Syn in the presence of 20% (w/v) PEG-8000 after 15 days of incubation at 25ºC. The inset represents associated α-Syn clusters in the same sample. (Right panel) Representative TEM image of the same sample showing the presence of amyloid fibrils. The experiment is performed twice with similar observations. **d.** Fluorescence microscopy images of 100 µM α-Syn in the presence of 15% (w/v) PEG-8000 spiked with 100 nM Alexa488-140C-α-Syn (top) and 50 µM ThT (bottom) at different timepoints (days 0-2) at 25ºC. The inset shows a magnified image of the ThT positive droplets at day 2. The experiment is performed 2 independent times with similar observations. **e.** (left panel) Normalized fluorescence recovery after photobleaching (FRAP) of α-Syn droplets (100 µM α-Syn+15% (w/v) PEG-8000) just after their appearance (day 1) is shown. Values represent mean ± S.D. for n=3 independent experiments. (Right panel) Representative snapshots of a α-Syn droplets at different times post-bleaching. The images are shown in thermal LUT for better visualization. **f.** Schematic describing two possible scenarios which explains delayed α-Syn assembly/LLPS from a thermodynamic perspective. In scenario (a), the crossing the nucleation energy barrier is the limiting step. On the other hand, in scenario (b), the growth of the assemblies is very slow. ΔG indicates Gibbs free energy.

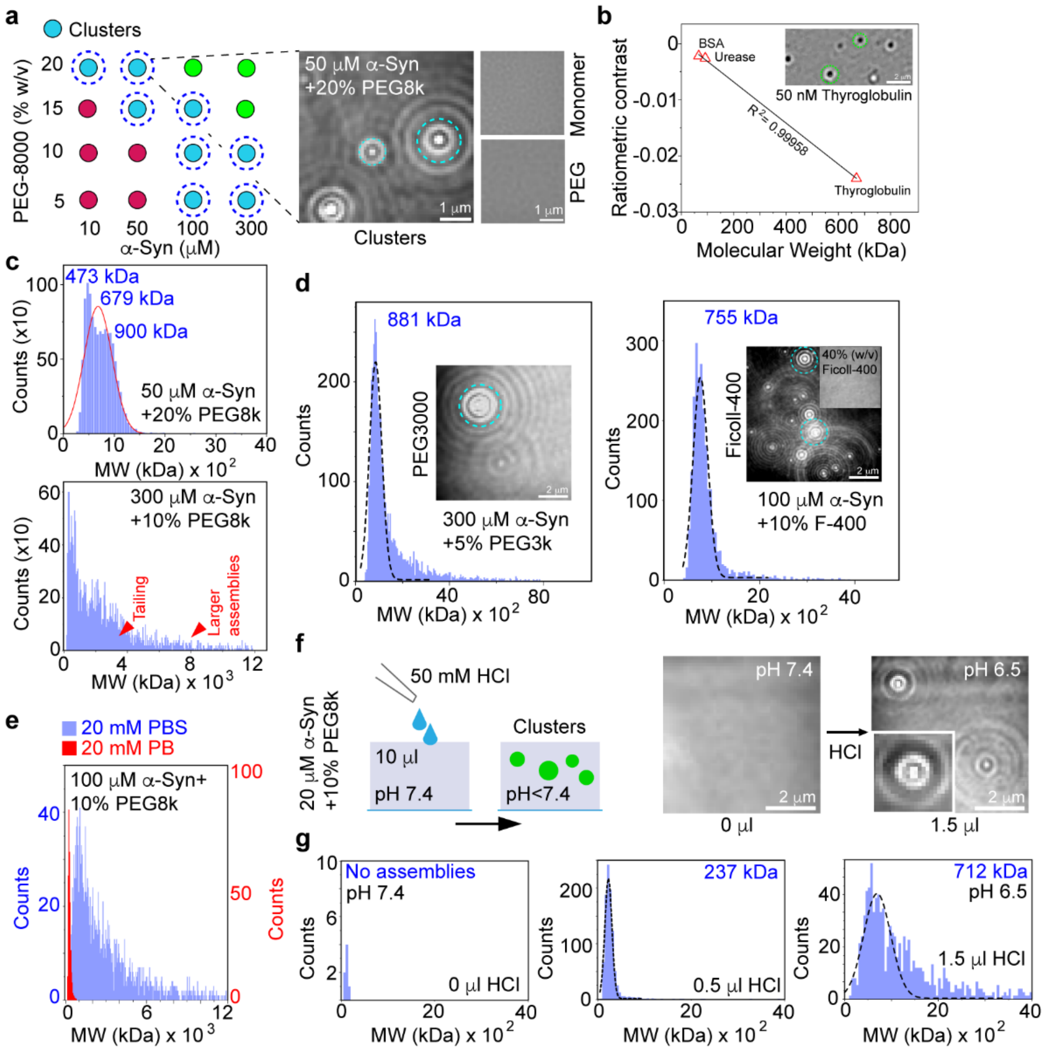


**Supplementary Figure 2. Mass photometry (MP) analysis of α-Syn nanoclusters: a.** (left panel) A schematic depicting the LLPS and nanocluster regime of α-Syn in presence of PEG-8000 at 25ºC. The green circles indicate canonical LLPS. The cyan circles indicate α-Syn nanoclusters which are visible by MP. The red circles indicate no assemblies. (Right panel) Representative MP image (native channel) of 50 μM α-Syn+20% (w/v) PEG-8000 showing presence of α-Syn clusters marked with cyan dashed circles. MP images (native channel) of only 550 μM α-Syn monomer and only 20% PEG-8000 (in 20 mM PBS, pH 7.4) are also shown. The experiments are performed 2 independent times with similar observation. **b.** A standard curve generated from measuring 50 nM BSA (66 kDa), urease (91 kDa, monomer) and thyroglobulin (669 kDa) is shown. The inset shows a MP image of 50 nM thyroglobulin monomers (669 kDa) as seen in the ratiometric channel. Individual events are marked with green dashed circles. **c.** Molecular weight (MW) histograms obtained from MP measurements of 50 μM α-Syn+20% (w/v) PEG-8000 (top) and 300 μM α-Syn+10% (w/v) PEG-8000 (bottom) are shown. The MW are noted in blue. For samples with multiple peaks (for example, 50 μM α-Syn+20% (w/v) PEG-8000, as shown here), the histogram is fitted to an in-built gaussian function (in red), and the average MW is noted (679 kDa). The red triangular pointers denote tailing of the histogram likely due to presence of larger assemblies. **d.** MW histograms obtained from MP measurements of 300 μM α-Syn+5% (w/v) PEG-3000 (left panel) and 100 μM α-Syn+10% (w/v) Ficoll-400 (right panel) are shown. The average MW is noted in blue. The insets show MP images of the corresponding samples and of 40% (w/v) Ficoll-400 (right) where no assemblies could be detected. **e.** MW histograms obtained from MP measurements of 100 μM α-Syn+10% (w/v) PEG-8000 prepared in 20 mM PBS (pH 7.4) (blue) and 20 mM PB (pH 7.4) (red) are shown. **f.** (left panel) Schematic describing the experimental procedure of pH titration in MP. Briefly, 10 μl of 20 μM α-Syn+10% (w/v) PEG-8000 (pH 7.4) is drop casted on the glass slide and 0.5 μl of 50 mM HCl is added to the solution in a stepwise manner. After each addition, the sample is measured in MP to check for α-Syn assembly. (Right panel) Representative MP images (native channel) of the sample before (left) and after (right) addition of 1.5 µl HCl are shown. g. MW histograms obtained from MP measurements of 20 µM α-Syn+10% (w/v) PEG-8000 (pH 7.4) before addition of HCl (left), and after addition of 0.5 µl (middle) and 1.5 µl HCl (right). The average MW is noted in blue. The experiment (f-g) is performed 2 independent times with similar observations.

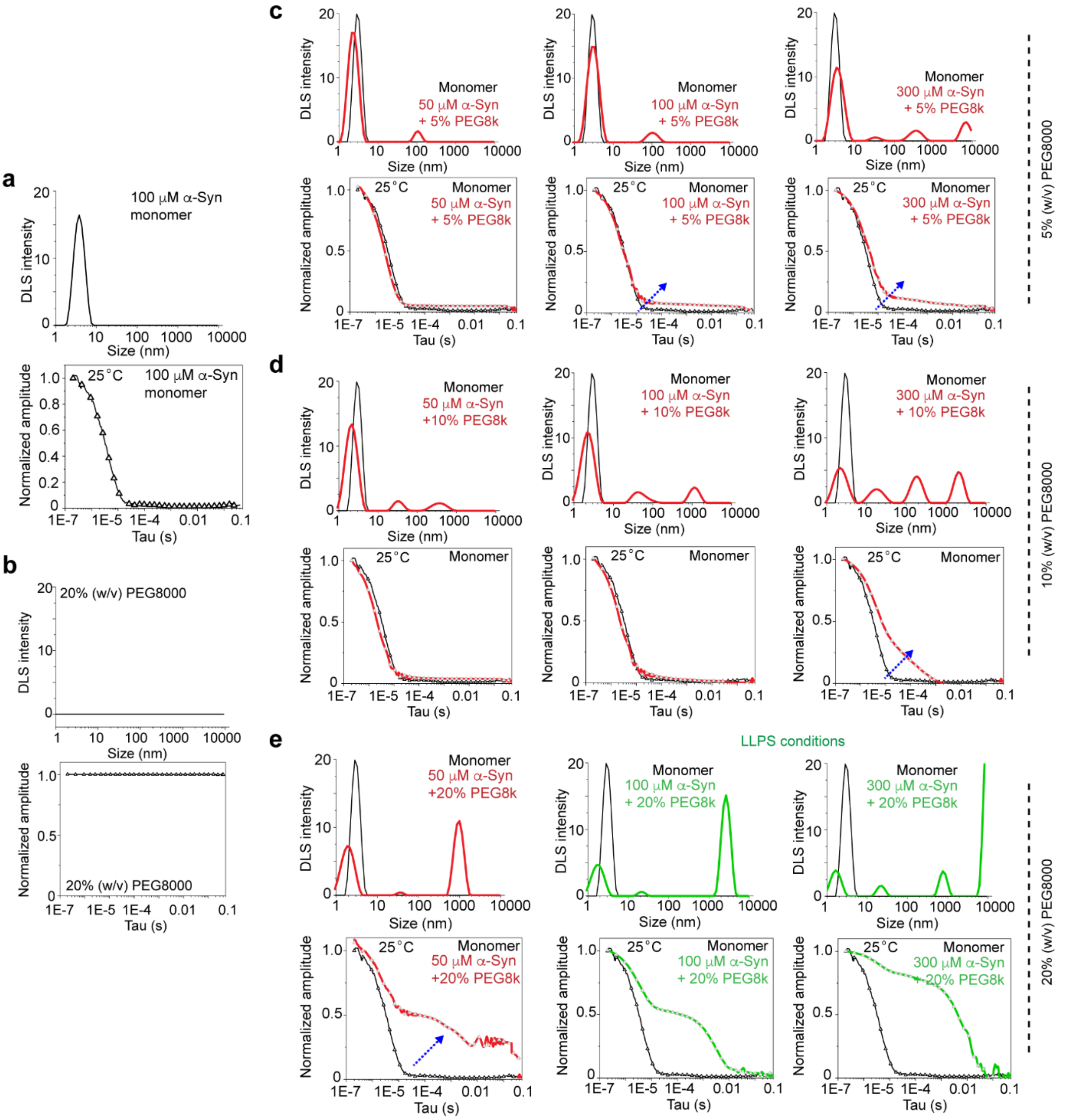


**Supplementary Figure 3. DLS studies to check for α-Syn assemblies below the apparent saturation concentration: a.** (upper panel) DLS profile of 100 μM α-Syn monomer showing a single peak at ~3.2 nm at 25ºC. The X-axis (size) is reported in log scale. (Lower panel) Normalized amplitude of the autocorrelation function of 100 μM α-Syn monomer is plotted. **b.** DLS intensity profile (upper panel) and normalized amplitude of the autocorrelation function (Lower panel) for 20% (w/v) PEG-8000 is shown. c-e. DLS intensity profile (upper panels) and normalized amplitude of the autocorrelation function (lower panels) for 50, 100 and 300 μM α-Syn in the presence of 5% (w/v) PEG-8000 (**c**); for 50, 100 and 300 μM α-Syn in presence of 10% (w/v) PEG-8000 (**d**); for 50, 100 and 300 μM α-Syn in presence of 20% (w/v) PEG-8000 (**e**) is shown. LLPS conditions are denoted in green. The blue arrows indicate the increase in the amplitude of the autocorrelation function for nanocluster forming samples. The data obtained for monomeric 100 μM α-Syn is also plotted as a reference in the same graphs (in black) for comparison (**c-e**). The X-axis are reported in log scale (**a-e**). All experiments are performed 3 times with similar results (**a-e**). All experiments are carried out at 25ºC.

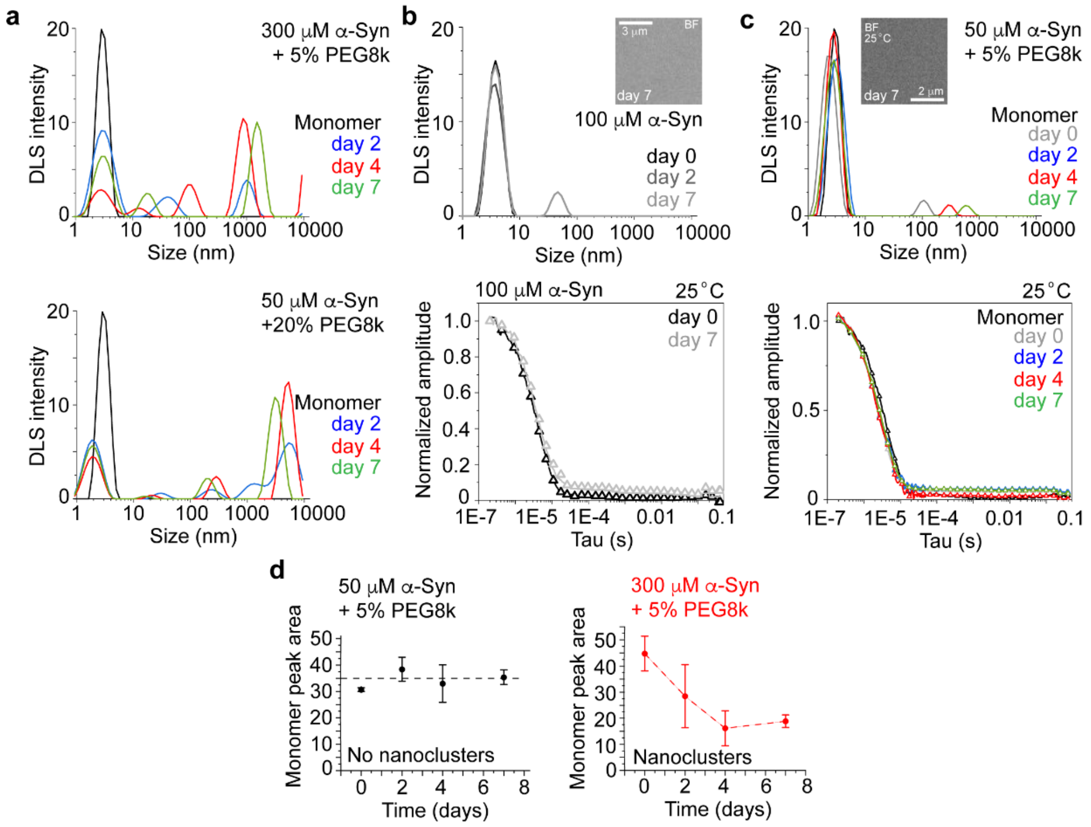


**Supplementary Figure 4. DLS measurements of α-Syn nanoclusters during their growth: a.** DLS intensity profile of 300 μM α-Syn+5% (w/v) PEG-8000 (top) and 50 μM α-Syn+20% (w/v) PEG-8000 (bottom) is shown for different timepoints (days 2-7) during incubation at 25ºC. Monomeric (100 μM) α-Syn is plotted as a reference in the same graphs (in black) for comparison. The experiments are carried out 2 independent times with similar results. **b-c.** DLS intensity profile (top) and normalized amplitude of the autocorrelation function (bottom) for monomeric (100 μM) α-Syn (**b**) and for 50 μM α-Syn+5% (w/v) PEG-8000 (**c**) is shown for days 0-7. The insets show representative BF microscopy images of the corresponding sample after 7 days of incubation at 25ºC. The experiments (**a-c**) are performed 3 times with similar results. **d.** The monomer peak area (1-10 nm) of 50 μM α-Syn+5% (w/v) PEG-8000 (no nanoclusters) (left) and 300 μM α-Syn+5% (w/v) PEG-8000 (nanoclusters) (right) is shown for days 0-7. The values represent mean ± S.D. for n=3 independent experiments.

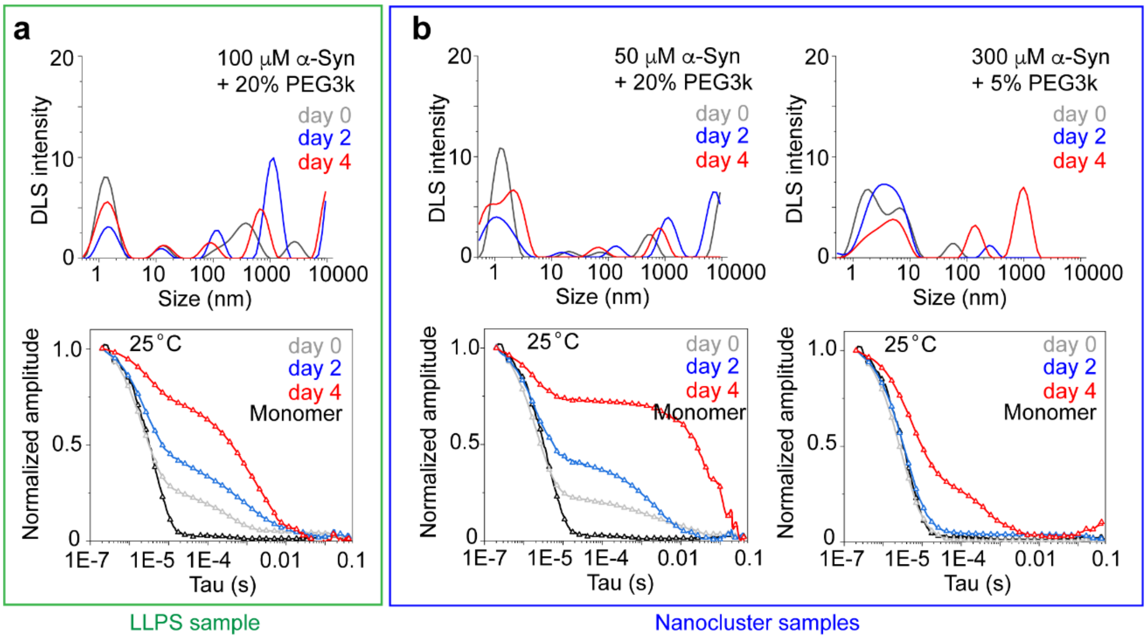


**Supplementary Figure 5. DLS measurements of α-Syn nanoclusters in presence of PEG-3000: a.** DLS intensity profile (top) and normalized amplitude of the autocorrelation function (bottom) for 100 µM α-Syn+20% (w/v) PEG-3000 (LLPS sample) is shown for days 0-4. **b.** (left panel) DLS intensity profile (top) and normalized amplitude of the autocorrelation function (bottom) for 50 µM α-Syn+20% (w/v) PEG-3000 (left panel) and for 300 µM α-Syn+5% (w/v) PEG-3000 (right panel) is shown for days 0-4. All experiments are carried out at 25ºC. The experiments (**a-b**) are performed 2 independent times with similar results.

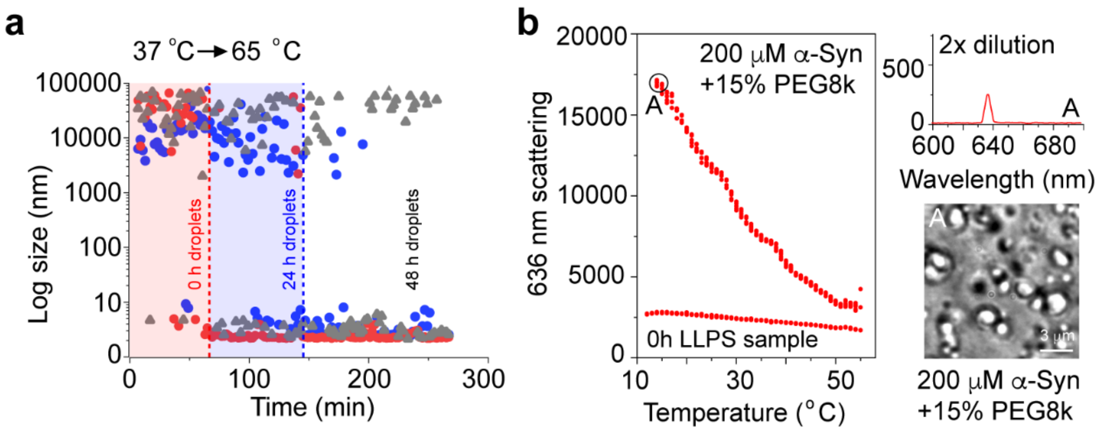


**Supplementary Figure 6. Effect of temperature on α-Syn clustering/LLPS: a.** Cumulant hydrodynamic radius of an LLPS sample, 100 µM α-Syn+20% (w/v) PEG-8000 (prepared in 20 mM PBS, pH 7.4) at different timepoints (0h, 24h and 48h) upon incubation at 65ºC for ~300 min. The data shows that newly formed α-Syn droplets take ~1h to dissolve. The dissolution time progressively increases when the LLPS sample is aged. **b.** (left panel) Static light scattering (SLS) intensity (636 nm) of 200 µM α-Syn+15% (w/v) PEG-8000 (prepared in 10 mM PB, 40 mM NaCl, pH 7.4) is shown while the sample is heated from 12ºC to 55ºC and back to 12ºC. (Right panel) After completion of the scattering experiment (at point A), the sample is diluted 2 times with buffer and the scattering intensity (636 nm) is plotted at 12ºC. The bottom panel shows a representative BF microscopy image of the droplets after completion of the SLS experiment (at point A). The experiments (a-b) are performed 2 times with similar results.

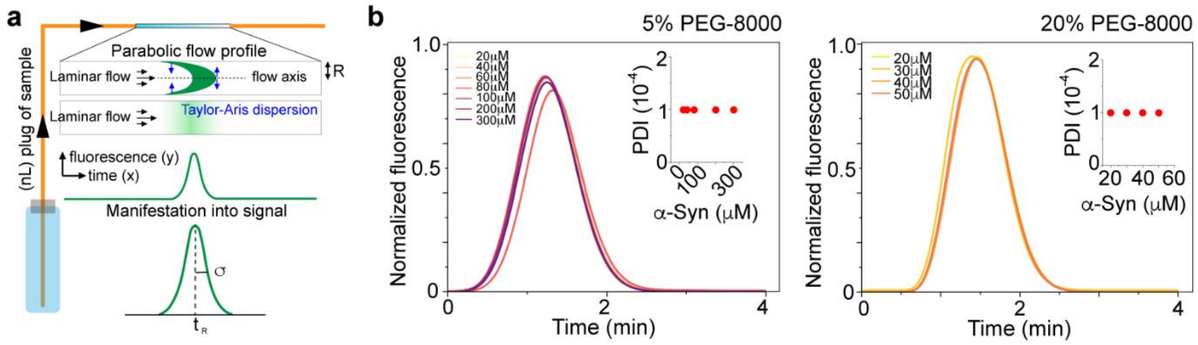


**Supplementary Figure 7. Flow induced dispersion analysis (FIDA) shows no polydispersity in nanocluster containing samples: a.** Schematic describing the FIDA[1] method. A plug of protein solution (~100 nl) is injected into a capillary having a radius R. Due to the laminar flow (different velocity at different distance from the center of the capillary), the protein plug gets distorted into a parabolic profile—eventually generating a gaussian gradient of distribution (Taylor-Aris dispersion). The time of arrival of this gaussian peak ($t_R$) and the variance ($\sigma$) helps us calculate the hydrodynamic radius and the PDI of the sample. **b.** Taylor grams[1] of 20-300 μM α-Syn in the presence of 5% (w/v) PEG-8000 (left) and 20-50 μM α-Syn in the presence of 20% (w/v) PEG-8000 (right) are shown. The insets show PDI values as a function of protein concentration. The experiment is performed twice with similar results.

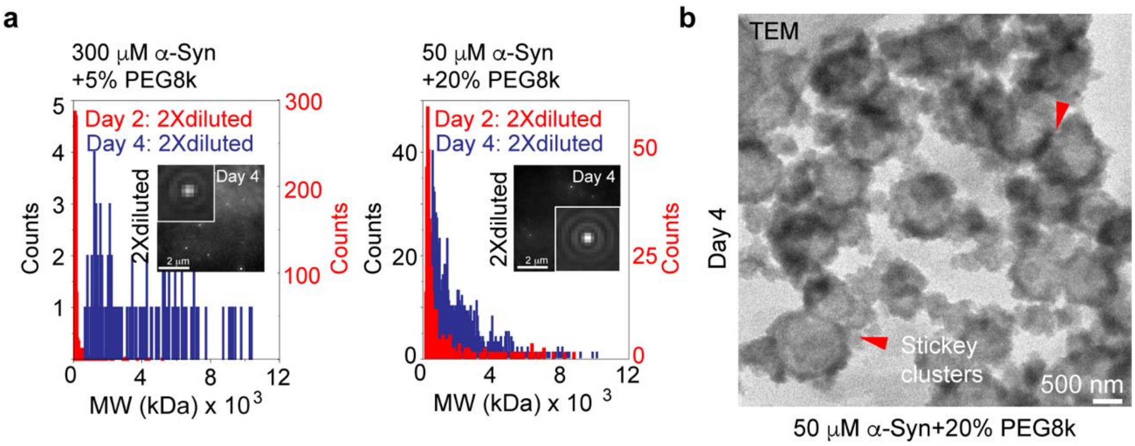


**Supplementary Figure 8. α-Syn nanoclusters solidify with time: a.** (Left panel) Molecular weight (MW) histograms obtained from MP measurements of 2X buffer-diluted samples: 300 µM α-Syn+5% (w/v) PEG-8000 (left) and 50 µM α-Syn+20% (w/v) PEG-8000 (right) are shown for day 2 (red) and day 4 (blue). Note that the large clusters persist even after dilution when the samples are aged for 4 days. Insets show representative MP image (native channel) of corresponding samples at day 4, showing the presence of α-Syn assemblies/clusters even after dilution. The experiments are performed 2 times with similar results. **b.** TEM image of 50 µM α-Syn+20% (w/v) PEG-8000 at day 4 showing presence of clumped α-Syn assemblies. The experiment is performed twice with similar results.

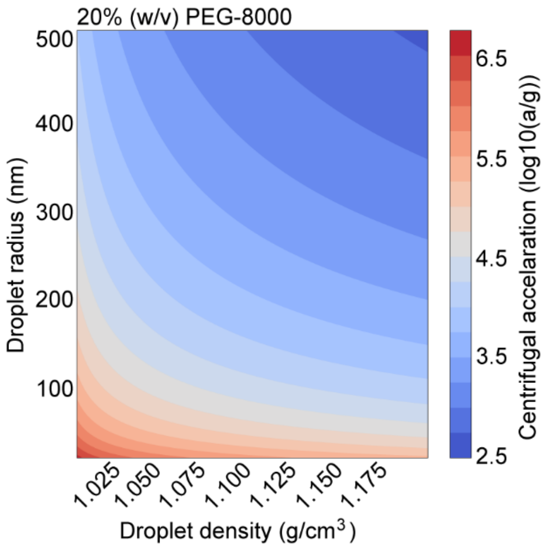


**Supplementary Figure 9. Centrifugal acceleration required to spin down protein assemblies in 20% (w/v) PEG-8000 solution:** A contour plot describing the centrifugal acceleration required to move spherical protein assemblies of different sizes (25-500 nm radius) and different densities (1.01-1.20 g/cm$^3$) by 1 cm in 1h in an aqueous solution containing 20% (w/v) PEG-8000 at 25°C. The color codes represent centrifugal acceleration (RCF) in log (10) scale.

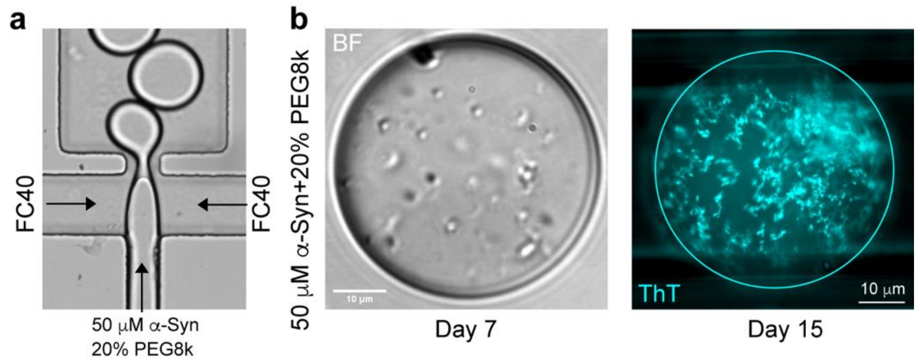


**Supplementary Figure 10. Delayed clustering of α-Syn in absence of surface or air-water interface: a.** BF microscopy image of the 3-component microfluidic channel showing encapsulation of 50 µM α-Syn+20% (w/v) PEG-8000 by FC-40. **b.** (left panel) BF microscopy image of a microdroplet after 7 days of incubation at 25ºC containing 50 µM α-Syn+20% (w/v) PEG-8000. (Right panel) Fluorescence microscopy image of a microdroplet containing 50 µM α-Syn+20% (w/v) PEG-8000 (spiked with 50 µM ThT) after 15 days of incubation at 25ºC. The experiments (**a-b**) are performed twice with similar observations.

**Supplementary movie 1. ThT fluorescence intensity from individual α-Syn droplets while ageing:** Movie showing emergence of ThT fluorescence from condensates formed by 200 µM α-Syn in presence of 20% (w/v) PEG-8000 in 20 mM PBS, pH 7.4 (spiked with 50 µM ThT) over a timescale of ~35h. The movie is captured with an interval of 20 min and at 800 ms exposure.

**Supplementary movie 2. Appearance of α-Syn clusters below saturation concentration:** Movie showing micron scale α-Syn clusters formed after 6 days of incubation at 25ºC under BF microscopy. 300 µM α-Syn in presence of 5% (w/v) PEG-8000 in 20 mM PBS, pH 7.4 is used for this experiment. The movie is captured with an interval of 100 ms.

**Supplementary movie 3. Delayed α-Syn assemblies are morphologically distinct:** Movie showing aggregation of 700 µM α-Syn in 20 mM PBS, pH 7.4 after 7 days of incubation at 25ºC under BF microscopy. The movie is captured with an interval of 10 min for ~10h.

**Supplementary movie 4. α-Syn nanoclusters in MP:** Movie showing α-Syn nanoclusters in MP for 50 µM α-Syn in presence of 20% (w/v) PEG-8000 in 20 mM PBS, pH 7.4 at 25ºC. The movie is captured above the coverslip surface for better visualization of the nanoclusters in the solution. The movie is captured in the native channel with an interval of 0.01 s for 1 min. No mass calculations are performed from this particular dataset.

**Supplementary movie 5. α-Syn nanoclusters close to the apparent saturation concentration ($C_{app}$):** Movie showing α-Syn nanoclusters in MP for 100 µM α-Syn in presence of 15% (w/v) PEG-8000 in 20 mM PBS, pH 7.4 at 25ºC. The movie is captured in the native channel with an interval of 0.01 s for 1 min. No mass calculations could be performed from this particular dataset because of detector saturation.

**Supplementary movie 6. Surface wetting by α-Syn nanoclusters as observed in MP:** Movie showing α-Syn nanoclusters in MP for 300 µM α-Syn in presence of 5% (w/v) PEG-8000 in 20 mM PBS, pH 7.4 at 25ºC. The movie is captured in the native channel with an interval of 0.01 s for 1 min. The data shows more and more nanoclusters landing on the coverslip surface with time.

**Supplementary movie 7. Delayed nucleation of α-Syn LLPS on a coverslip surface:** Movie showing many α-Syn droplets nucleating after ~4-5h in BF microscopy for 100 µM α-Syn in presence of 20% (w/v) PEG-8000 in 20 mM PBS, pH 7.4 at 25ºC. The movie is captured with an interval of 10 min for ~17h.

**Supplementary movie 8. α-Syn nanoclusters can form without surface nucleation:** Movie showing α-Syn assemblies forming inside a FC-40 coated microdroplet after 7 days of incubation at 25ºC. A standard nanocluster forming condition, 50 µM α-Syn in presence of 20% (w/v) PEG-8000 in 20 mM PBS, pH 7.4, is used for this experiment. The movie is captured with an interval of 8 s for ~15 min.
